# Prospects of telomere-to-telomere assembly in barley: analysis of sequence gaps in the MorexV3 reference genome

**DOI:** 10.1101/2021.11.08.467736

**Authors:** Pavla Navrátilová, Helena Toegelová, Zuzana Tulpová, Yi-Tzu Kuo, Nils Stein, Jaroslav Doležel, Andreas Houben, Hana Šimková, Martin Mascher

**Author notes:** These authors contributed equally (Pavla Navrátilová, Helena Toegelová). Correspondence should be addressed to Hana Šimková or Martin Mascher.

## Abstract

The first gapless, telomere-to-telomere (T2T) sequence assemblies of plant chromosomes were reported recently. However, sequence assemblies of most plant genomes remain fragmented. Only recent breakthroughs in accurate long-read sequencing have made it possible to achieve highly contiguous sequence assemblies with a few tens of contigs per chromosome, i.e. a number small enough to allow for a systematic inquiry into the causes of the remaining sequence gaps and the approaches and resources needed to close them. Here, we analyze sequence gaps in the current reference genome sequence of barley cv. Morex (MorexV3). Optical map and sequence raw data, complemented by ChIP-seq data for centromeric histone variant CENH3, were used to estimate the abundance of centromeric, ribosomal DNA and subtelomeric repeats in the barley genome. These estimates were compared with copy numbers in the MorexV3 pseudomolecule sequence. We found that almost all centromeric sequences and 45S ribosomal DNA repeat arrays were absent from the MorexV3 pseudomolecules and that the majority of sequence gaps can be attributed to assembly breakdown in long stretches of satellite repeats. However, missing sequences cannot fully account for the difference between assembly size and flow cytometric genome size estimates. We discuss the prospects of gap closure with ultra-long sequence reads.

## Introduction

The recent advances in genome sequencing and assembly methodology have enabled the gap-less reconstructing of the sequences of entire eukaryotic chromosomes. Telomere-to-telomere (T2T) assemblies have been reported for one maize chromosome (Liu et al., 2020), five banana chromosomes (Belser et al., 2021) and the human genome (Logsdon et al., 2021; Miga et al., 2020; Nurk et al., 2021). T2T assembly requires the use of complementary sequence and mapping resources for primary contig assembly, scaffolding and gap closure. For example, Logsdon et al. (2021) used a combination of accurate long reads (PacBio HiFi) and ultra-long nanopore reads for sequence assembly and, among other resources, a Bionano optical map and manually curated sequences of bacterial artificial chromosome (BAC) clones for validation. These resources were generated and analyzed with the express purpose of closing all remaining gaps in the sequence of human chromosome 8, with a particular focus on completing the sequence of centromeric satellite repeat arrays. By contrast, reference genome projects in crops did not focus on T2T sequence but rather aimed at near-complete gene space representation, chromosome-scale contiguity at the scaffold level, and reasonable representation of the repeat space. The expenditure and effort required to achieve these goals has decreased drastically in recent years as sequence reads became longer and more accurate and powerful scaffolding methods such as optical genome mapping and chromosome-conformation capture sequencing (Hi-C) were developed. Even for plant species with large, heterozygous or autopolyploid genomes, near-complete chromosome-scale sequence assembly has become possible (Sun et al., 2021; Zhang et al., 2019; Zhou et al., 2020).

Whether it is possible and how much effort it will take to turn a “near-complete” into a finished sequence assembly of a plant genome have become spertinent questions. The report of Liu et al. (2020) illustrates that T2T assembly is not impossible, but also not easy: even with a combination of long-read technologies and optical mapping, the centromeric sequence of only a single maize chromosome could be completed. As of today, all plant reference genome sequences, even that of *Arabidopsis thaliana*, have gaps (Naish et al., 2021). An assessment of what is missing in current plant reference genomes is a timely undertaking. Before embarking on the quest for a T2T assembly, it should be ascertained as best as possible with the resources at hand which challenges will have to be overcome and which additional datasets are needed to achieve gapless assemblies of entire chromosomes. The genome of barley (*Hordeum vulgare* L.) is a good model to study the prospects of T2T assembly in plant because (i) it has a high-quality reference sequence assembly and (ii) much is known about its repeats. Mascher et al. (2021) used accurate long-read sequencing (PacBio HiFi), Bionano optical mapping and chromosome-conformation capture sequencing (Hi-C) to construct the latest barley reference sequence (MorexV3), which surpasses the older versions MorexV1 (Mascher et al., 2017) and MorexV2 (Monat et al., 2019) in gene space completeness and the representation of recently inserted transposable elements (Mascher et al. 2021). Accurate long reads spanning entire elements of the predominant retrotransposon families have reduced the number of sequence gaps in the pseudomolecules from hundreds of thousands in the MorexV1 and MorexV2 short-read assemblies to only 147, of which 51 were spanned by contigs of a Bionano optical map.

A meaningful assessment of missing sequence in near-complete genome assemblies requires knowledge about the repeat composition that is independent of sequence assemblies. Over the past decades, scientists have studied the extent and arrangement of barley telomeres (Kilian et al., 1995; Röder et al., 1993), subtelomeres (Belostotsky and Ananiev, 1990a; Brandes et al., 1995; Röder et al., 1993), centromeres (Houben et al., 2007; Hudakova et al., 2001; Presting et al., 1998) and ribosomal DNA (rDNA) loci (Leitch and Heslop-Harrison, 1992; Leitch and Heslop-Harrison, 1993) using cytological and molecular biology techniques. Large and homogeneous repeat arrays constituting these functional loci are prime candidates for difficult-to-assemble sequences missing from the current barley reference sequence assembly as suggested by our previous analysis of subtelomeric and 45S rDNA repeats (Kapustová et al. 2019). Here, we conduct an in-depth analysis of the representation of repeat arrays in the latest reference sequence assembly of barley cv. Morex and its underlying raw data. Our results indicate functional genomic loci of barley such as centromeres and ribosomal DNA clusters can be assembled only if and when sequence reads longer than 100 kb or even 1 Mb become available.

## Results

We based our analyses on the current reference genome sequence assembly of barley cv. Morex (MorexV3, Mascher et al. (2021)) and several publicly available short-read and long-read datasets **(Table 1**; **Supplementary Figures 1-3)**, a genome-wide Bionano optical map and newly generated chromatin immunoprecipitation sequencing (ChIP-seq) data for centromere histone H3 (CENH3). Using these data, we compared estimates of genome size from different sequence datasets with those from flow cytometry and quantified missing sequences at (sub-) telomeric satellites, centromeric repeats, ribosomal DNA and other satellite repeats.

**Table 1:** Summary statistics and genome size estimates (GSE) from different sequence datasets.

|  | HiFi<sup>1</sup> | ONT<sup>1</sup> | PE450<sup>1</sup> |
| --- | --- | --- | --- |
| Number of reads | 6.6 M | 32.5 M | 684.6 M |
| Sequenced base pairs | 132.7 G | 426.9 G | 364.2 G |
| Proportion of aligned reads | 100.0% | 98.9% | 98.5% |
| Mean depth | 31.4 | 94.3 | 81.8 |
| Median depth | 31 | 85.2 | 76.2 |
| Mode depth | 31.3 | 84.3 | 76.4 |
| GSE based on read depth (Gb)<sup>4</sup> | 4.3 | 5.0 | 4.8 |
| GSE based on 21-mer spectra (Gb) | 2.9 | 0.3 | 3.8 |
| GSE based on 51-mer spectra (Gb) | 4.2 | - | 4.3 |
| GSE based on 101-mer spectra (Gb) | 4.3 | - | - |

<sup>1</sup>HiFi: PacBio HiFi reads (accurate long reads); ONT Oxford Nanopore long reads; PE450 2x250 bp Illumina reads of fragments with an average size of 450 bp.
<sup>4</sup>The genome size estimate was calculated as the number of sequenced base pairs divided by the median depth.

### Discrepancy between genome size estimates

A simple approach to assess the completeness of a genome sequence assembly is to compare the size of the assembly with the estimated size of the whole genome. To date, the only DNA sequence-independent methods that have been used to estimate genome size in plants are Feulgen microspectrophotometry and flow cytometry. Due to the ease of use and higher throughput, flow cytometry gradually replaced the former approach (Doležel and Bartoš, 2005). Flow cytometric estimations of haploid (1C) DNA amounts of *H. vulgare* ranged from 3.64 pg (Marie and Brown, 1993) to 5.47 pg (Vaikonen, 1994), corresponding to 3.559 Gb - 5.349 Gb. Both microspectrophotometric and flow cytometric assays need a reference standard with known genome size (Doležel and Bartoš, 2005) and as there is currently no plant or animal species with known genome size that is suitable as a reference standard, all published data are based on genome sizes arbitrarily assigned to the standards. This is one of the reasons why the estimates for the same species may differ between studies (Doležel and Greilhuber, 2010). Recently, Doležel et al. (2018) estimated 1C genome size of barley cv. Morex as 4.88 Gb and 5.04 Gb, respectively, using human leukocytes as the primary reference standard, considering two different values for the human genome size.

The seven pseudomolecules of the MorexV3 assembly amount to 4.196 Gb, with 29.1 Mb in unplaced contigs. The preceding versions MorexV1 (Mascher et al., 2017) and MorexV2 (Monat et al., 2019) were longer than MorexV3, with assembly sizes of 4.834 Gb and 4.343 Gb, respectively. However, both MorexV1 and MorexV2 suffered from limitations of short-read assembly that have led to overestimated assembly sizes. MorexV1 harbored redundant sequences because of incomplete merging of fragmented sequence assemblies of overlapping bacterial artificial chromosome (BAC) clones. In MorexV2, gap sizes in scaffolds constructed from mate-pair reads may have been mis-estimated (Monat et al., 2019). The assembly size of MorexV3 may be lower than the true genome size because accurate long reads may suffer from sequence dropout in long low-complexity G/A- or T/C-rich regions (Nurk et al., 2020).

Assembly-free genome size estimates (GSEs) are commonly obtained by evaluating *k*-mer frequency spectra computed from high-throughput sequencing data. The original method proposed by Li and Waterman (2003) has been refined by tools such as GenomeScope (Vurture et al., 2017) and findGSE (Sun et al., 2018). We used findGSE with different *k*-mer sizes (21, 51, 101) on (1) PacBio HiFi circular consensus reads (HiFi); (2) Oxford Nanopore (ONT) reads and (3) paired-end short reads (2×250bp, PCR-free) from 450 bp fragments (PE450). The HiFi and ONT datasets were by Mascher et al. (2021); the PE450 data by Monat et al. (2019). At *k*=51, HiFi and PE450 yielded concordant results with GSEs in the range of 4.2 Gb and 4.3 Gb, respectively **(Table 1**). At *k*=21, estimates for both datasets were lower, indicating that GSEs in barley are influenced by the choice of *k*-mer size. ONT reads gave no meaningful estimate at *k*=21; the use of a larger *k* was not feasible due to the high error rate.

An alternative method to infer genome sizes is based on the average coverage in read alignments to assemblies (Pfenninger et al., 2021): 10 Gb of shotgun reads from a 1 Gb genome will result in an average 10-fold read depth. When this argument is reversed, we can infer from a 10x average coverage for 10 Gb of shotgun reads a genome size of 1 Gb. We aligned HiFi, PE450, and ONT reads to the MorexV3 pseudomolecules and summarized read depths and calculated GSEs **(Table 1**). Estimates from short reads and accurate long reads were in the range of those derived from *k*-mer spectra and similar to the MorexV3 assembly size. The estimate from the uncorrected ONT reads (5 Gb) was close to the higher flow cytometric estimate (5.04 Gb) by Doležel and Greilhuber (2010). However, we found that 9.3 % of the ONT reads aligned to the barley chloroplast (cp) genome for ≥ 90 % of their length. ONT reads matching to the cp genome amounted to 29.5 Gb, which, if truly originating from the nuclear genome, would be equivalent to 347 Mb of plastid DNA insertions amounting to about half a barley chromosome. However, fluorescence *in situ* hybridization (FISH) with a probe specific for barley cpDNA did not support the presence of large (> 100 kb) cpDNA insertions into the nuclear genome of barley cv. Morex: only rather weak hybridization signals, about as strong as those of labelled cDNA clones (Aliyeva-Schnorr et al., 2016), were seen on chromosomes 2H and 5H (**Fig. 1**). Both chromosomes were identified based on the distribution of the 45S rDNA and subtelomeric satellite repeat HvT01 (Szakács and Molnár-Láng, 2007). Hence, the DNA preparations for ONT sequencing were likely contaminated with cpDNA. After introducing a correction factor into our coverage calculation, we arrived at a coverage-based genome size estimate of 4.7 Gb.

**Figure 1:**
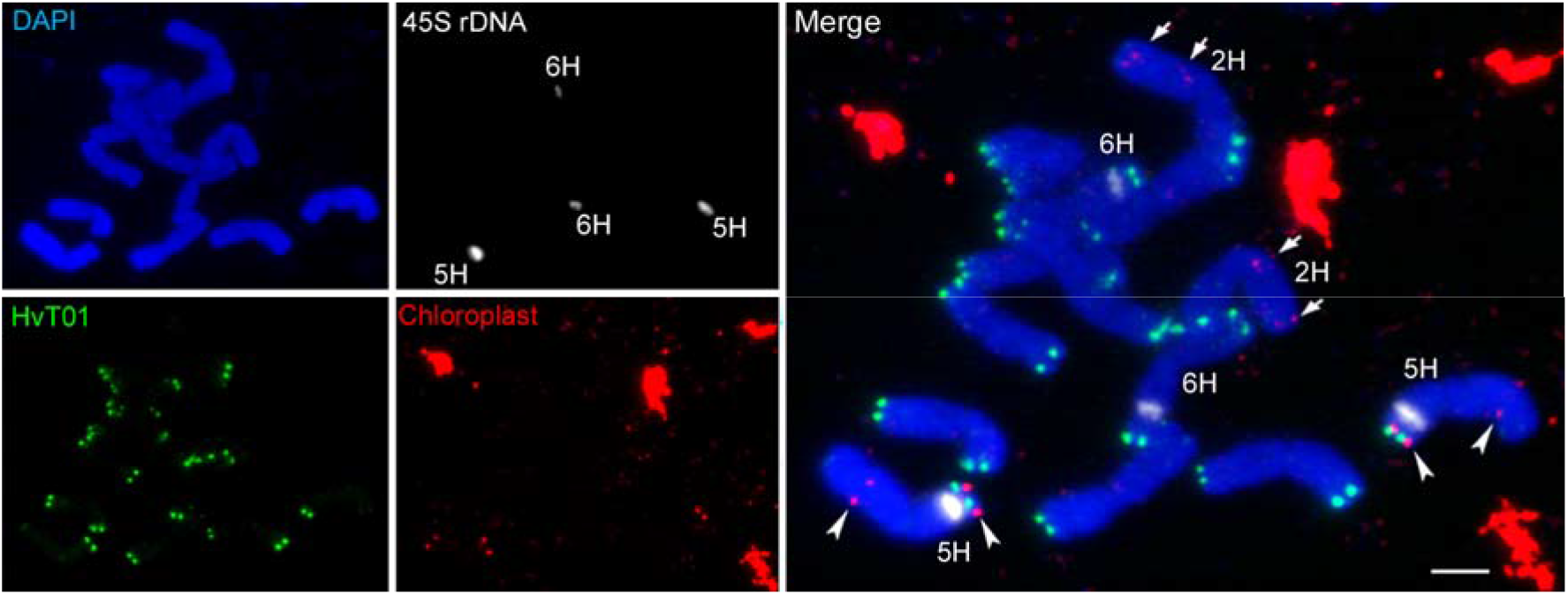
FISH mapping of chloroplast insertions into the nuclear genome of barley cv. Morex. Chloroplast DNA (red), subtelomeric satellite HvT01 (green) and 45S rDNA (white) were mapped on metaphase chromosomes. Chloroplast derived DNA insertions are detected on chromosome 2H (*arrows*) and 5H (*arrowheads*). Non-chromosomal chloroplast DNA signals represent plastids. Chromosomes were counterstained by DAPI. Scale bar = 5 μm.

Previous studies have reported discrepancies between flow cytometric and sequence-based GSEs, although the reasons have remained unclear (Pflug et al., 2020). In the following, we analyze candidates for difficult-to-assemble loci to understand if gaps in the genome assembly can account for the large difference (up to hundreds of Mb or 10 % of the genome) between various flow cytometric estimates and MorexV3 assembly size.

### Telomeric satellite arrays are not captured in their entirety by long reads

Telomeres of barley chromosomes consist of thousands of TTTAGGG repeat copies (Kilian et al., 1995). To assess the completeness of the MorexV3 pseudomolecules (Mascher et al., 2021) at the chromosomal ends, we aligned the pseudomolecules to the Morex Bionano optical map, which revealed missing sequences at the termini of all short arms and of three long arms **(Table 2, Fig. 2)**. The missing segments at short arm termini were generally longer (17-220 kb) than those in the long arms (10-80 kb). It is to be noted that the truly missing sequence at the chromosomal ends is larger than these estimates since the optical map is likely incomplete in the terminal regions because DLE-1 recognition sites, required for labelling molecules in optical mapping, are absent from telomeric and most subtelomeric repeats. Search for the TTTAGGG motif in the pseudomolecules revealed continuous arrays of 3.6-13.7 kb at three long-arm termini and discontinuous arrays interspersed by other sequences at the ends of two additional arms. Interestingly, all partially assembled arrays of telomeric repeats were on long arms while all short arms missed relatively large segments at their termini. A similar trend was observed in the long-read B73 RefGen v4 assembly of maize (Jiao et al., 2017) where a part of a telomeric array has been assembled in seven out of ten long arm ends but only two of the short arms. A possible explanation are differences in the copy number and homogeneity of subtelomeric repeats between long and short arms.

**Table 2.** Completeness of MorexV3 assembly in telomeric regions.

| Chromosome | Optical map overhang<sup>1</sup><br>(kb) |  | Telomeric repeat<sup>2</sup><br>in sequence<sup>2</sup> (kb) |  |
| --- | --- | --- | --- | --- |
|  | Short arm | Long arm | Short arm | Long arm |
| 1H | 100 | - | - | + |
| 2H | 140 | 10 | - | 3.6 |
| 3H | 220 | - | - | 13.7 |
| 4H | 154 | - | - | + |
| 5H | 20 | 10 | - | 5 |
| 6H | 70 | 80 | - | - |
| 7H | 17 | - | - | - |

<sup>1</sup>Terminal part of the optical map extending beyond the start/end of a pseudomolecule, indicating a missing sequence.
<sup>2</sup>Presence of an interspersed (subtelomeric) array of TTTAGGG repeats is marked by +. Lengths are given for regular telomeric arrays only.

**Figure 2.**
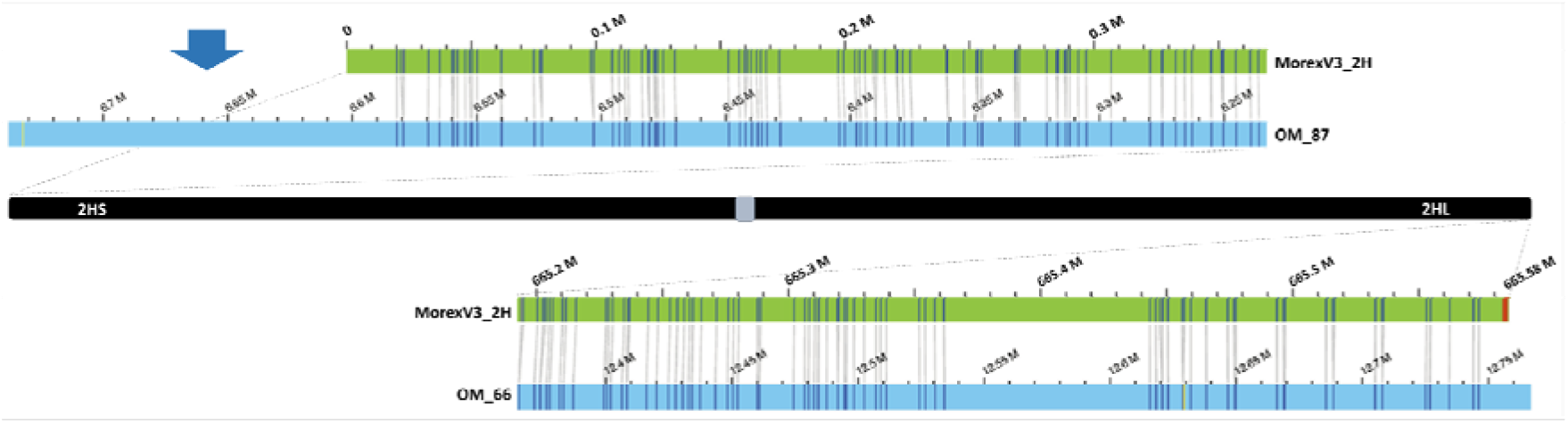
Alignments of the termini of the 2H optical map and sequence assembly reveal missing sequence. The optical map (blue bar) was aligned to the 2H pseudomolecule sequence (green bar). Vertical grey lines connect matching DLE-1 label sites (CTTAAG motif). The optical map extends beyond the 2H sequence at the short arm (2HS) terminus (blue arrow), indicating a minimum of 140 kb missing sequence, while at the long arm (2HL) terminus, the sequence contains 3.6 kb of a regular telomeric motif (TTTAGGG, marked by red stripes) belonging to a functional telomere. Long label-free map segments at both termini suggest the presence of unlabelled subtelomeric satellite repeats.

To complement this analysis, we investigated telomeric satellite repeats in the PacBio HiFi and ONT long reads that were used to construct the MorexV3 pseudomolecules (Mascher et al., 2021). Tandem Repeat Finder (TRF) was used to annotate arrays of tandem repeats and satellites on all individual read sequences. We found that ONT reads with TTTAGGG arrays longer than 1 kb were mapped to distal ends of only three chromosome arms (2HL, 3HL, 5HL, **Fig. 3a)**. TTTAGGG arrays were also found in the distal 2 Mb of 3HS and 7HL, but not at the very end of the pseudomolecule sequence, indicating either the presence of interstitial arrays or errors in sequence orientation. More than half of reads (51 %) with TTTAGGG arrays > 1 kb matched to unanchored contigs. Arrays longer than 1 (10) kb were found in a total of 1848 (452) ONT reads. The average size of TTTAGGG arrays >= 1kb was 6.9 kb; the longest TTTAGGG array annotated in an ONT read spanned 37.2 kb. As this read was entirely composed of TTTAGGG motifs, the length of the complete array it belongs to is unknown. The cumulative size of TTTAGGG arrays longer than 1 kb annotated on the ONT reads was 13.0 Mb. Assuming an average 85-fold coverage with ONT reads, this amounts to an average of 11 kb of telomeric satellites per chromosome arm. This is shorter, but on the same order of magnitude, as the telomere size of ~22 kb estimated by terminal restriction fragment analysis (Kilian et al., 1995). We found only 44 TTTAGGG arrays longer than 1 kb in HiFi reads with a cumulative length of 317 kb, amounting to 741 bp of non-redundant sequence per telomere (assuming 31x genome coverage). This clear underestimate may be explicable by HiFi sequence dropout in GA-rich regions (Nurk et al., 2020).

**Figure 3:**
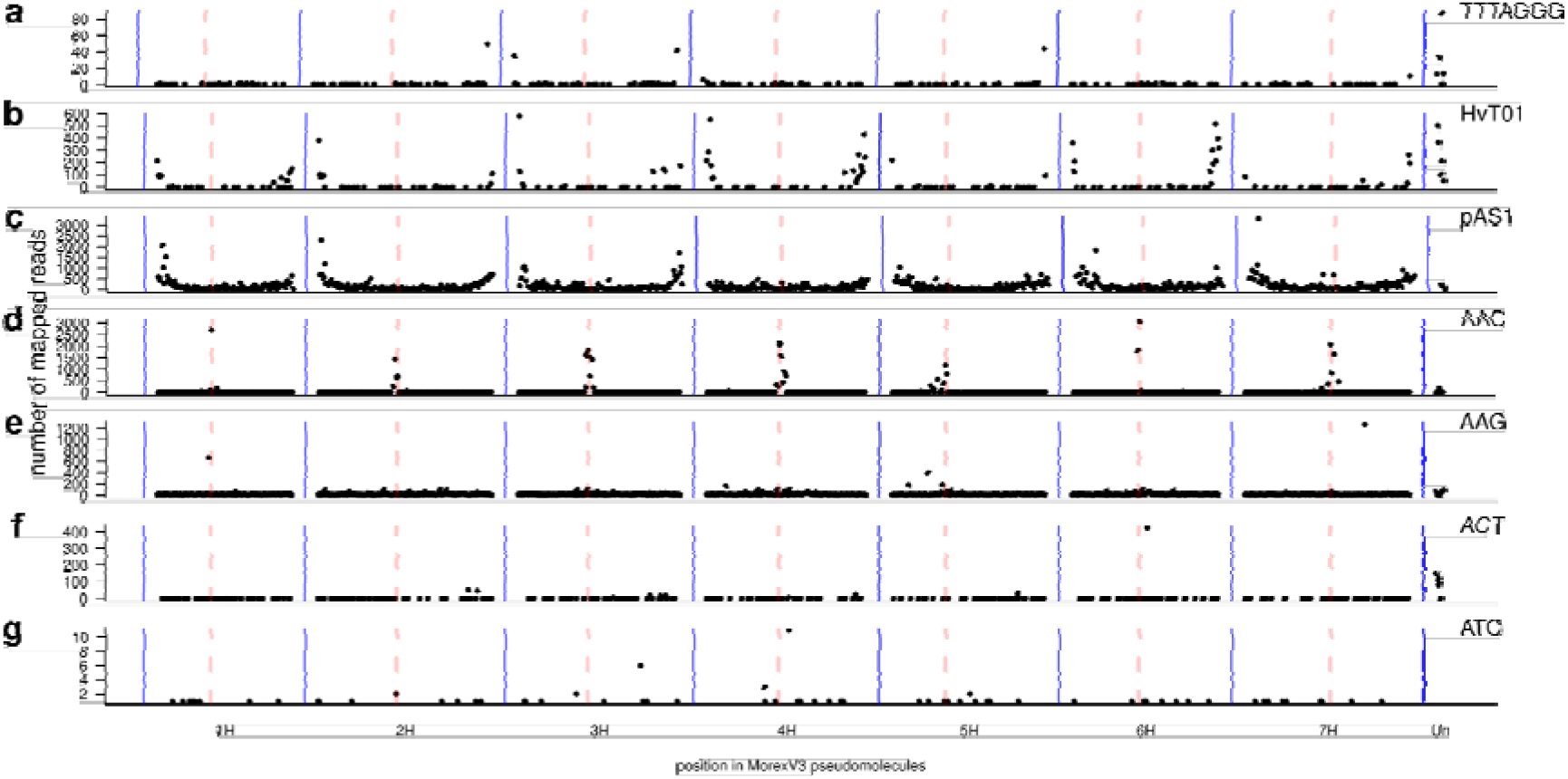
Locations of known satellite arrays in the MorexV3 pseudomolecules. Known repeat sequences were annotated in read sequences. Telomeric repeats (TTTAGGG, **(a)**) and trinucleotide microsatellites (**d-g**) were identified with Tandem Repeat Finder in ONT reads; subtelomeric repeats [HvT01 **(b)** and pAS1 **(c)]** were found by BLAST against HiFi reads. The counts of reads containing these repeats and mapped uniquely to positions in the MorexV3 pseudomolecules were aggregated in 5 Mb windows and plotted along the genome. Dashed red lines mark centromeres.

### Subtelomeric repeat arrays are disrupted by sequence gaps

To assess the representation of subtelomeric satellite repeats in the MorexV3 assembly, we studied two previously reported repeats, HvT01 and pAS1. The 118 bp subtelomeric repeat HvT01 was first reported by Belostotsky and Ananiev (1990a) and was mapped by FISH to the distal ends of all chromosome arms of barley (Schubert et al., 1998; Szakács and Molnár-Láng, 2007). Brandes et al. (1995) discovered pAS1, a 336 bp sequence with a preferentially subtelomeric localization.

We aligned the HvT01 and pAS1 consensus sequences to the HiFi reads using BLAST (Altschul et al., 1990) and selected near-complete alignments (>= 110 bp for HvT01; >= 330 bp for pAS1). The mean length of HvT01 alignments was 20 kb, i.e. spanning entire reads. This indicates that HvT01 is present in long arrays. By contrast, the length of pAS1 alignments was 6.6 kb on average. Plotting the alignment positions of reads positive for either HvT01 or pAS1 indicated that HvT01 is only found in very distal locations, whereas pAS1 is present throughout the chromosomes, albeit with a strong enrichment towards the distal ends **(Fig. 3b,c)**.

We focused on reads that had alignments to either HvT01 or pAS1 with cumulative lengths of 1 kb or more. A total of 3,803 reads satisfied this criterion for HvT01, amounting to a cumulative alignment length of 97.6 Mb. Assuming 31-fold coverage with HiFi reads, the total size of HvT01 arrays is estimated at 3.1 Mb, i.e. on average 225 kb of HvT01 sequence per chromosome arm. A total of 158,607 reads contained long (> 1 kb) alignments to pAS1 with a cumulative length of 804 Mb, equivalent to a non-redundant sequence of 25.9 Mb (1.85 Mb per chromosome arm). We attempted to use also ONT reads for an independent estimation of subtelomeric repeat abundance, but observed a 5-10 fold difference in cumulative alignment length between identity cut-offs of 70 and 80 %, indicating that a high degree of sequence divergence relative to the consensus monomer prevents confident alignment to uncorrected long-reads.

Alignments of the consensus monomers of HvT01 and pAS1 to the MorexV3 assembly covered 3.7 Mb and 28.2 Mb of sequence, respectively – largely consistent with our estimates based on read depth. Only 134 kb (0.5 %) of the pAS1 sequence were matched to unanchored contigs (“chrUn”), while a substantial fraction (19.3 %, 707 kb) of HvT01 were assigned to chrUn. The likely reason is that long stretches of homogeneous HvT01 arrays led to ambiguities in the assembly graph, resulting in short contigs that could not be assigned to chromosomal locations.

### Functional centromeres are absent from the pseudomolecules

Independent of their underlying DNA sequences, functional centromeres of most species are defined by the presence of the centromere-specific histone H3 variant CENH3 (Talbert and Henikoff, 2020). The two main constituents of barley centromeres are the *Ty3/gypsy*-retrotransposon *cereba* and the AGAGGG satellite repeat (Hudakova et al., 2001; Presting et al., 1998). Chromatin immunoprecipitation (ChIP) for CENH3 showed that both *cereba* and AGAGGG satellites interact with CENH3 (Houben et al., 2007). We attempted to position centromeres in the MorexV3 sequence assembly without using prior knowledge of centromeric sequences, following two complementary approaches: the inspection of contact probability matrices and ChIP-sequencing (ChIP-seq). Contact probabilities were determined from chromosome conformation capture sequencing (Hi-C) data of cv. Morex (Mascher et al., 2017) and visualized as two-dimensional matrices recording the number of Hi-C links between pairs of genomic loci (Lieberman-Aiden et al., 2009). Contact matrices of all barley chromosomes showed a characteristic cross pattern with a strong main diagonal and a weaker anti-diagonal **(Fig. 4)**. We had previously interpreted this pattern as a consequence of the Rabl configuration of interphase nuclei (Mascher et al., 2017). The Rabl configuration refers to a clustering of centromeres and telomeres of all chromosomes during interphase, juxtaposing loci from opposite arms at the same relative distance from the centromere (Cowan et al., 2001; Rabl, 1885). As an alternative visualization of Hi-C data, we used directionality indices recording imbalances in the number of upstream and downstream contacts along the genome (Dixon et al., 2012; Himmelbach et al., 2018). The midpoints of the “Rabl crosses” coincided with strong discontinuities in the directionality indices **(Fig. 4)**, which we assume corresponded to the locations of functional centromeres. To support the assertion that this discontinuity marks the position of the centromeres, we conducted ChIP-seq for barley α-CENH3 (Ishii et al., 2015; Sanei et al., 2011) to determine the locations of DNA sequences bound to centromeric nucleosomes. The observed CENH3 ChIP-seq peaks fell together with the jump in the directionality index and also colocalized with AGAGGG arrays in all chromosomes **(Fig. 5a, c, d; Supplementary Figs. 4-9)**. For most chromosomes, several additional CENH3 peaks were observed in the pericentromeric region. Both the satellite array and the ChIP-seq peaks were narrow, spanning tens of kilobases at the most (cumulatively less than 200 kb for all peaks at a single centromere), and there were only 17-54 of *cereba* retroelements in the 10 Mb regions around the directionality breakpoints. This is at odds with previous estimates of about 200 *cereba* elements per barley centromere (Presting et al., 1998), corresponding to at least 1.4 Mb of sequence (Hudakova et al., 2001). The small width of the ChIP-seq peaks also contrasts to analogous results in bread wheat (*Triticum aestivum*). In this species, CENH3 ChIP-seq peaks occupy intervals of several megabases on all chromosomes (The International Wheat Genome Sequencing Consortium (IWGSC), 2018). All major ChIP-seq peaks in barley were adjacent to gaps between contigs in the pseudomolecules, spanning sequences of unknown size. None of the gaps was bridged by contigs of the optical map **(Supplementary Fig. 10)**, indicating the presence of long (> 100 kb) stretches of DNA devoid of DLE-1 label sites. Nevertheless, non-aligned and sparsely labelled map overhangs of tens to more than a hundred kilobases, extending into the assumed centromeric gaps of some chromosomes support the presence of missing sequence in the pseudomolecules **(Supplementary Fig. 10)**. We did not find any ONT reads spanning a centromere gap, supporting the notion that centromeres are longer than 100 kb.

**Figure 4:**
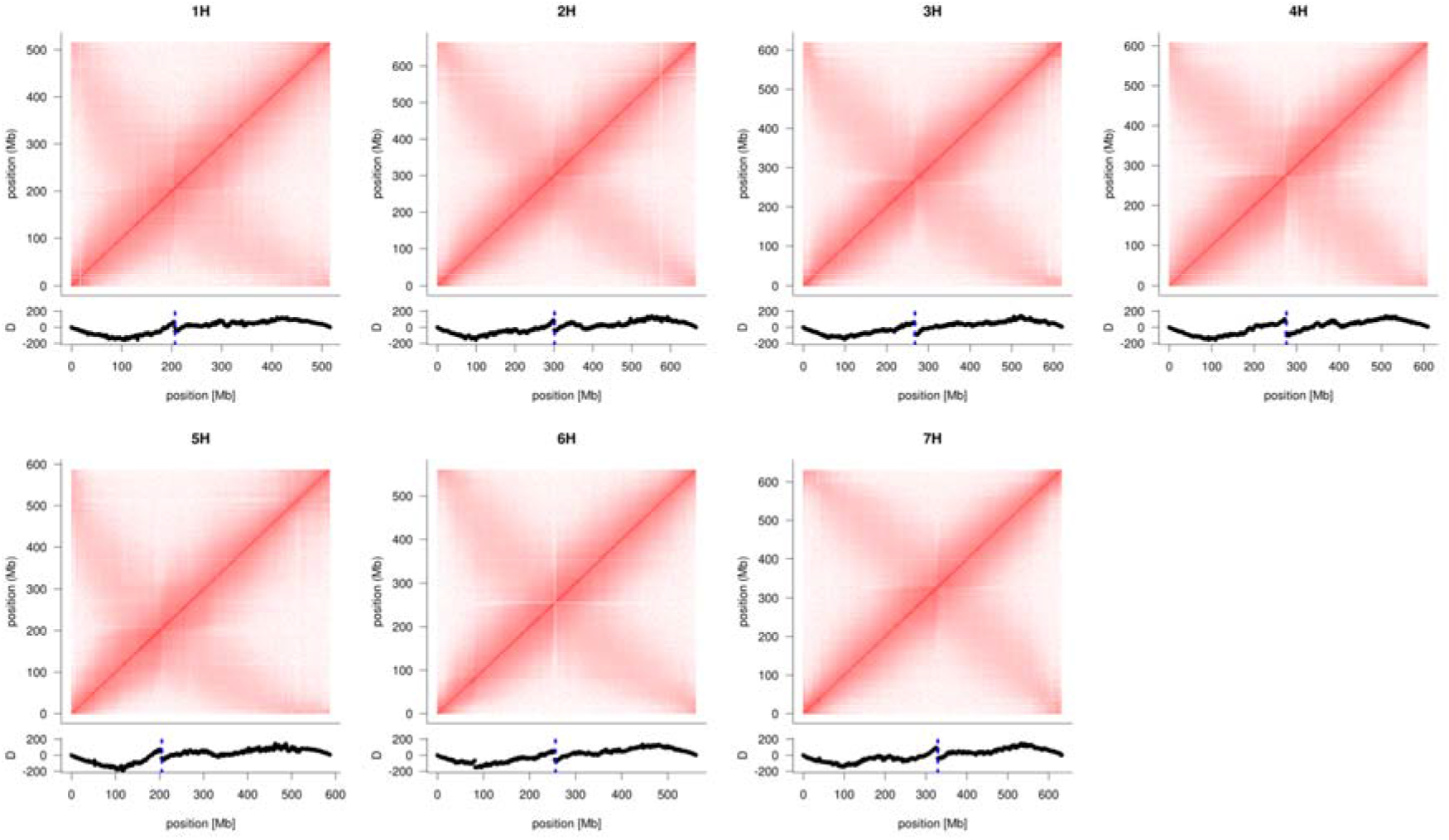
Positioning of centromeres by inspection of Hi-C contact matrices. Intrachromosomal Hi-C contact matrices for the seven barley chromosomes were computed from alignment of Hi-C reads to the MorexV3 pseudomolecules. The intensity of the red color is proportional to the contact probability. Below each contact matrix is shown the directionality bias (excess of up- or downstream Hi-C links) along the chromosomes. Discontinuities coinciding with the intersection points of the diagonals and anti-diagonals mark putative centromeres locations (marked by blue dotted line).

**Figure 5:**
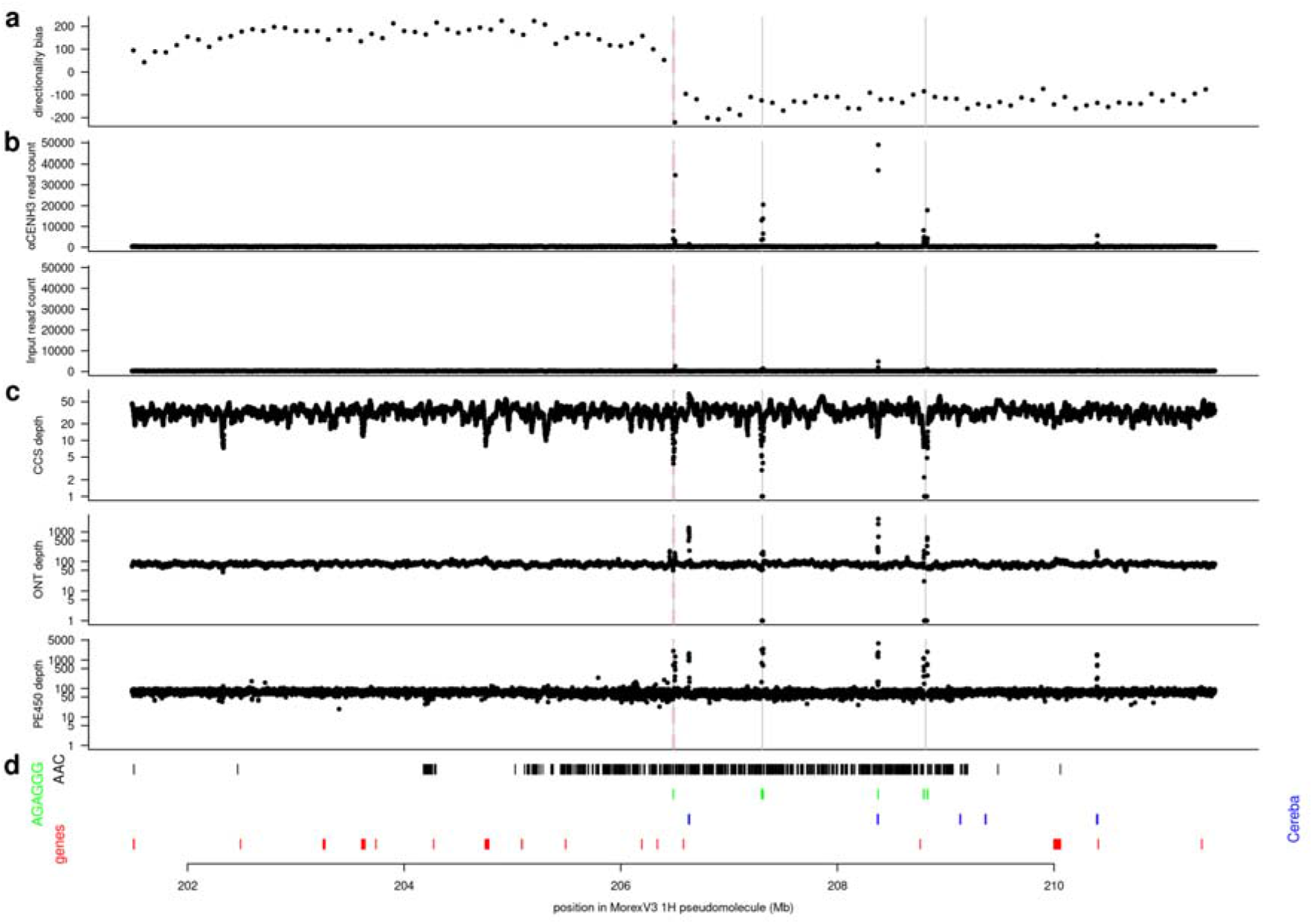
Sequence organization of the pericentromeric region of chromosome 1H. A 10 Mb region flanking (+/- 5 Mb) a gap likely containing the functional centromere are shown. In panels **(a)-(c)**, the centromere is indicated by a dashed red line; gaps in the pseudomolecules are marked by vertical gray lines. **(a)** Directionality bias in non-overlapping 100 kb windows. **(b)** Read depth in CENH3 ChIP-seq data and the input controls. Data from two biological replicates were summed. **(c)** Read depth in HiFi, ONT and PE450 data. **(d)** Position of (i) AAC (black) and AGAGGG (green) satellite arrays; (ii) Cereba elements (blue); (iii) genes (red). Analogous plots for chromosome 2H to 7H are shown in **Supplementary Figs. 4-9**.

As sequences interacting with CENH3 may be missing from the pseudomolecules or their repetitive nature may prevent unambiguous mapping of ChIP-seq reads, we analyzed the α-CENH3 ChIP-seq data in a reference-free manner using RepeatExplorer2 followed by ChIP-seq Mapper tool (Novák et al., 2020). RepeatExplorer2 uses graph-based clustering of the whole-genome shotgun (WGS) reads for characterization of repetitive DNA, resulting in a set of repeat clusters. ChIP-seq Mapper further assigns the ChIP and input reads to those repeat clusters and reports ChIP/input ratios of the normalized read counts for each cluster. The most strongly enriched sequence cluster (CL78, α-CENH3-ChIP-seq/input ratio = 11) was composed of a mix of *cereba* sequences and the AGAGGG-type of satellite repeats and was estimated to occupy 0.43 % of the barley genome, i.e. 20.21 Mb, assuming a genome size of 4.7 Gb **(Supplementary Fig. 11)**.

As an alternative approach to estimating the number of centromeric repeats, we ran TRF and BLAST alignments on the long reads of cv. Morex as we did for (sub-)telomeric repeats. The 1.4 Mb size estimate of Hudakova et al. (2001) is based on the copy number of the *cereba* integrase domain given by Presting et al. (1998). The latter authors had extrapolated from phage library screens for integrase that barley chromosomes contain on average 200 *cereba* copies, each 7 kb in size. We performed BLAST searches with the 809 bp integrase domain against the HiFi reads. A total of 19,090 (25,471) reads contained a near-complete (≥800 bp alignment length) hit at 90 % (80 %) sequence identity. Ninety-eight per cent of these reads had only a single hit, indicating that the barley lambda 9 clone of Hudakova et al. (2001) with two *cereba* copies in close proximity was an exception. Assuming a 31-fold coverage, the number of *cereba* integrase copies amounts to ~100 per chromosome – an estimate on the same order of magnitude, albeit somewhat lower, than the 200 copies of Presting et al. (1998).

The notion that individual *cereba* copies are separated by intervening satellite arrays was supported by the TRF results for the AGAGGG satellite. A total of 3,159 HiFi reads contained a satellite array longer than 1 kb. Of these, only 13.2 % were mapped with high confidence (MAPQ = 60) to positions more than 1 Mb away from a centromere gap, 69 % were unmapped and 17.4 % mapped within 1 Mb of a centromere gap. These mapping results are concordant with the presence of FISH signals for AGAGGG only at the centromeres (Houben et al., 2007; Hudakova et al., 2001; Kapusi et al., 2012). Among the 3,159 AGAGGG-positive reads, 35.6 % contained a *cereba* integrase. BLAST alignments of the 7 kb sequence of a complete *cereba* element showed that only 19 % of integrase-positive reads had a full-length hit to *cereba*. However, partial hits (>= 2000 bp alignment length) were found for 93 %, indicating that *cereba* elements may be rapidly disrupted after insertion.

The results for ONT reads were largely consistent with those for HiFi reads. A total of 86,708 ONT reads had near-complete integrase hits at 80 % identity, which corresponds to 146 copies per chromosome (assuming 85-fold coverage). Long (>= 1 kb) AGAGGG arrays were found in 67,481 ONT reads. Of the AGAGGG-positive ONT reads, only 18 % mapped more than 1 Mb away from the centromere gaps and 23 % had a BLAST alignment to the *cereba* integrase. The longest AGAGGG array found in the ONT reads spanned 95 kb; arrays longer than 30 kb were found in 346 ONT reads. These large array sizes explain the gaps found at the centromeres in the MorexV3 pseudomolecules, which were constructed from HiFi reads selected for smaller size ranges (15-22 kb) and possibly defective in their coverage of A/G rich motifs (Nurk et al., 2020). We note that while ONT reads with AGAGGG arrays longer than 1 kb amount to 2.3 Mb of non-redundant sequence, AGAGGG-positive HiFi reads can account only for 245 kb of non-redundant sequence, an observation possibly related to HiFi sequence dropout in GA-rich regions (Nurk et al., 2020).

A rough estimate for the average centromere sizes of barley chromosomes based on the *cereba* copy number and the *cereba* to AGAGGG ratio is as follows: 100 copies of mostly incomplete *cereba* elements with an average length of 4 kb amount to 400 kb. Alignments to *cereba* accounted for 11.5 % of the sequence of HiFi reads containing an AGAGGG array longer than 1 kb. Assuming that *cereba* elements make up one eighth of a functional centromere, the average centromere size is 3.2 Mb, similar to the estimate derived from the analysis of repeats in ChIP-seq data (20.21 Mb / 7 = 2.89 Mb). Most of this sequence is missing from the MorexV3 pseudomolecules.

### Size estimation of ribosomal DNA by optical maps and accurate long reads

One of the most important functional domains in the nucleus is the nucleolar organizer region. It consists of 45S ribosomal DNA, which is arranged in long arrays of homogenous units composed of clusters of highly conserved 18S, 5.8S and 26S rRNA genes separated by intergenic spacers, whose size and sequence composition can differ between particular loci in a genome **(Fig. 6a)**. To date, ribosomal DNA arrays are not present in most genome assemblies, including the most recent human reference genome GRCh38.p13 (Schneider et al., 2017).

**Figure 6:**
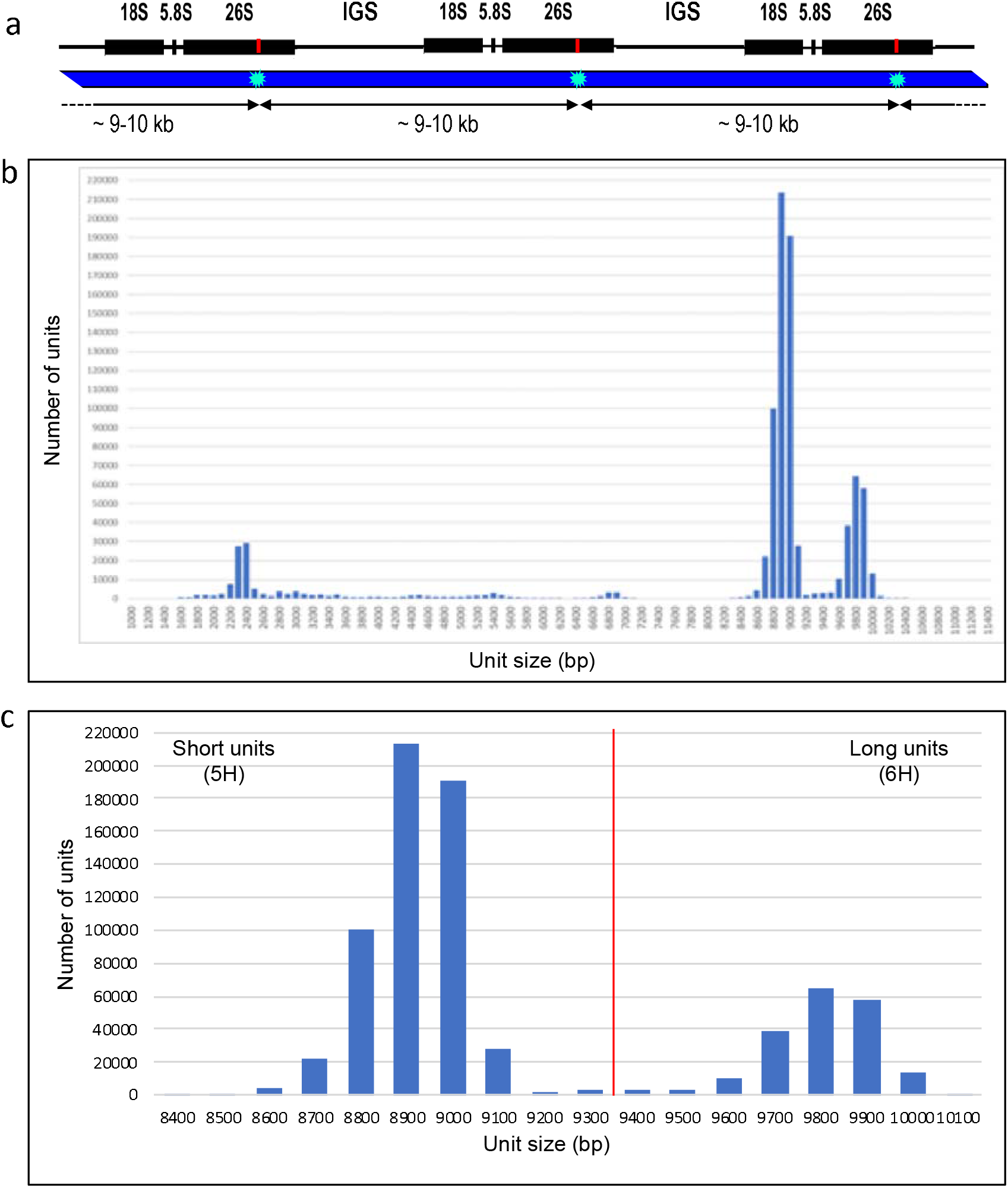
Profiling of 45S rDNA tandem repeats in optical map data. **(a)** 45S rDNA arrays are composed of clusters of 18S, 5.8S and 26S rRNA genes separated by intergenic spacers (IGS) whose lengths differ between 5H (total unit size ~8.9 kb) and 6H (total unit size ~9.8 kb) rDNA loci. Labelling at DLE-1 site in the 26S rRNA gene (red line) generates a regular pattern in the optical map with a label spacing of ~9-10 kb (bottom). **(b)** Size distribution of DLE-1-labelled tandem repeats in Morex optical map raw data >150 kb, considering repeat arrays >5 units. The dataset totals 1.09 Tb corresponding to 237-fold coverage of the barley genome. **(c)** The size category of 8,400-10,100 bp belongs to 45S ribosomal DNA.

We previously proposed Bionano optical mapping as a valuable tool to position and characterize particular rDNA loci (Tulpová et al., 2021) and assess their completeness in reference genomes (Kapustová et al., 2019). To figure out what portion of the 45S rDNA had been included in the reference genome, we exploited raw data used to generate the DLE-1 optical map. Whole-genome profiling of DLE-1-labelled arrays with more than five units revealed three major size categories of labelled tandem repeats – 2.2-2.5 kb, 8.6-9.1 kb and 9.6-10 kb **(Fig. 6b)**. The sizes of the latter two corresponded to the sizes of 5H and 6H rDNA units, respectively, identified by BLAST searches in the interval 52.6-53.7 Mb on the 5H and 81.9-82.4 Mb on the 6H pseudomolecule of MorexV3. These positions correspond to the major 45S rDNA loci in Morex barley **(Fig. 1**). Alignment of the 5H and 6H pseudomolecules to the DLE-1 optical map (Mascher et al., 2021) showed maps with the expected regular pattern aligned to the identified rDNA positions, but none of the maps spanned across the whole rDNA region **(Supplementary Fig. 12)**. Manual inspection of the Bionano map contigs showed that the regular pattern with ~9-10-kb spacing occurred only in those aligning to the rDNA positions or in several shorter unassigned contigs that did not comprise other pattern and were distinguished by high molecule coverage. We conclude that all Bionano molecules with the ~9-10 kb pattern most likely belonged to rDNA arrays and that the spacing of 8.6-9.1 and 9.6-10 kb corresponded to 5H and 6H units, respectively **(Fig. 6c)**. Analysis of a dataset totalling 1.09 Tb (232-fold coverage of the 4.7 Gb genome of Morex) assigned a total of 5.042 Gb (565,420 units) and 1.886 Gb (192,345 units) to the short and long rDNA units, respectively. We estimate that the 5H and 6H loci comprised 2,435 and 829 regularly arranged rDNA units, respectively, corresponding to 21.71 and 8.12 Mb of sequence for the 5H and 6H loci, respectively. BLAST searches in the 5H pseudomolecule of MorexV3 identified rDNA arrays of 102 and 48 complete units in 5H and 6H, respectively, positioned in the interval 52.6-53.7 Mb.

An additional 11.42 Mb of both complete and incomplete 45S rDNA units were found in unassigned scaffolds (chrUn) **(Supplementary Fig. 12)**. Based on the cumulative size of these alignments and the optical map-based rDNA abundance estimate of 29.8 Mb (0.64 % of the Morex genome), at least 16 Mb of rDNA sequence are missing in the MorexV3 assembly.

To confirm the results from Bionano genome mapping, we estimated the abundance of 45S rDNA in our long-read data. Representative sequences of the 45S unit on chromosomes 5H and 6H, respectively, were aligned to the HiFi reads. Considering alignment longer than 5 kb with at least 90 % sequence identity, 41,228 reads with a cumulative length of 820 Mb were aligned to both the 5H and 6H unit. Only 16 reads (amounting to 213 kb) were aligned to only one of the units. Of the reads aligned to both units, 2,480 (49.8 Mb, 6 %) were assigned to locations in the MorexV3 pseudomolecules with mapping quality ≥ 10, mainly to chromosomes 5H, 6H and 1H. These reads may correspond to degenerate copies close to the boundaries of the major rDNA arrays on 5H and 6H and to the degenerated minor locus on 1H. Assuming that (i) 95 % (787 Mb) of reads with hits to both units are intact sequences originating from one of the major arrays on 5H and 6H and that (ii) the average HiFi read depth is 31 **(Supplementary Fig. 1**), we arrive at an estimated size of 25.4 Mb of both arrays combined. This sequence-based estimate is 15 % smaller than the one based on the Bionano map.

To assess the representation of 5S rDNA, we aligned its 120 bp coding sequence (Fukui et al., 1994) to the HiFi reads and the MorexV3 pseudomolecules **(Fig. 7)**. We found 9,971 hits with > 90 % identity and > 100 bp alignment. Of these, 6,586 were on chr2H in the interval 575-577 Mb. Smaller arrays were found on chromosomes 4H (573 Mb, 256 copies) and 7H (250 Mb, 967 copies). Isolated hits (< 20 copies) were reported on chromosomes 1H, 3H and 5H. A large number (2138 copies) were on unassigned contigs. A total of 321,943 hits to the HiFi reads were reported, equivalent to ~10,400 unique copies at 31-fold coverage. This indicates that the majority of 5S rDNA gene copies are represented in the MorexV3 assembly, although ~21 % of them are on contigs not assigned to a chromosomal location.

**Figure 7:**
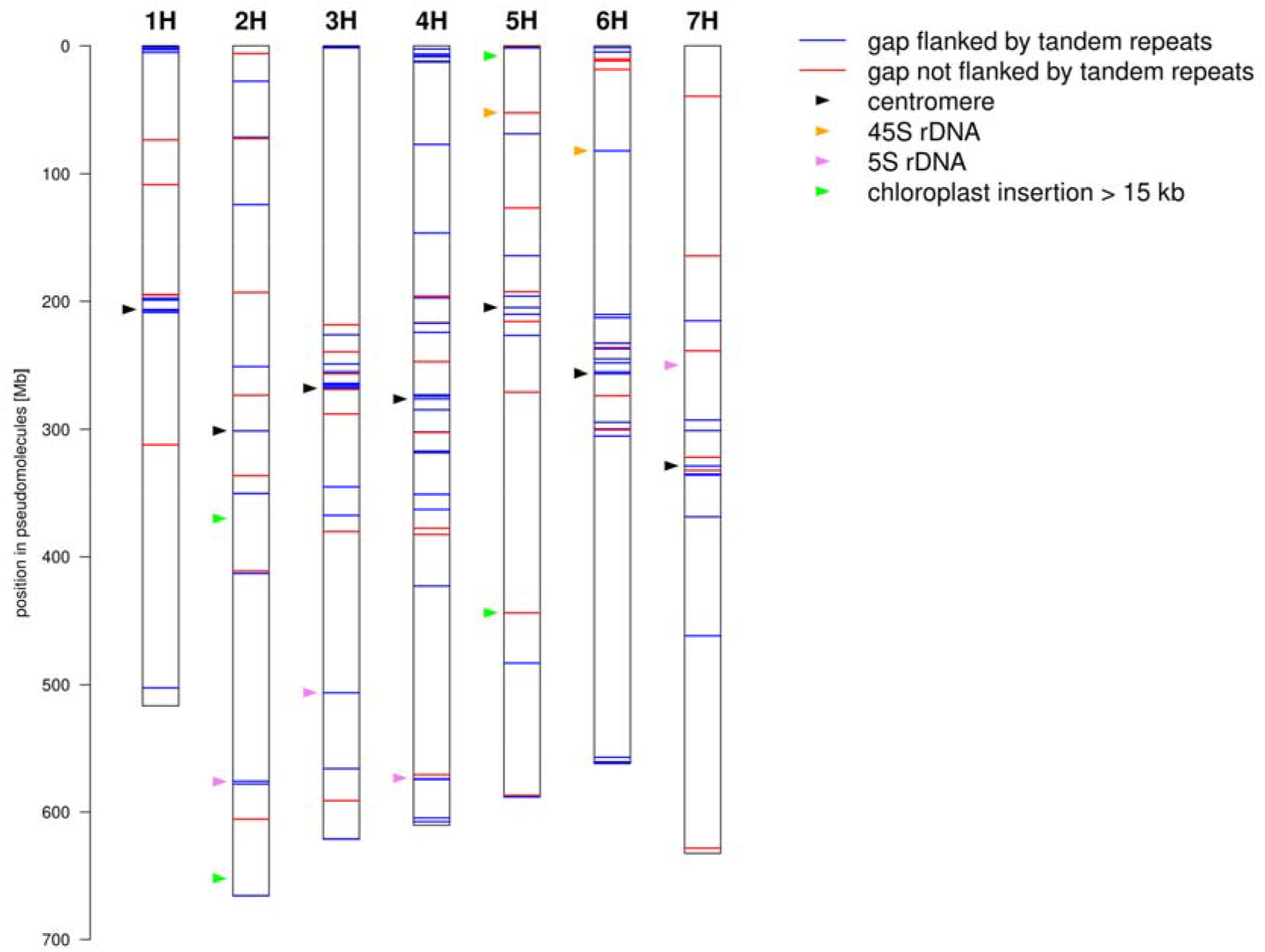
Sequence gaps in the MorexV3 pseudomolecules. The positions of sequence gaps are shown along the pseudomolecules. Gaps are colored according to whether or not at least one of their 1 kb flanking regions contain tandem repeat arrays longer than 500 bp. The positions of centromeres, 5S and 45S ribosomal DNA loci and large (> 15 kb) insertions of plastidal DNA are marked by colored arrowheads.

### Most sequence gaps are due to repeat arrays

Our analyses so far have shown long, homogeneous satellite repeat arrays are represented only incompletely in the MorexV3 pseudomolecules. We asked ourselves how many other sequence gaps can be attributed to long stretches of low-complexity DNA. TRF identifies all repeat arrays with a maximum motif size of 2 kb. We inspected ONT reads for the presence of long (> 20 kb) repeat arrays, which cannot be spanned by HiFi reads. Longer ONT reads may span entire gaps or at least enable the positioning of long arrays by the presence of a single-copy sequence at least at one end of a read. Among the most abundant motifs identified by TRF in the ONT reads were the trinucleotide microsatellites AAC, AAG, ACT and ATC. A total of 7,655 reads contained trinucleotide arrays longer than 20 kb. The longest array with 50,473 AAG copies spanned 153 kb. Trinucleotide microsatellites had been mapped using FISH by Cuadrado and Jouve (2007). Unique sequences adjacent to repeat arrays made it possible to assign repeat-containing ONT reads to chromosomal locations **(Fig. 3d-g)**. Consistent with the previous FISH mapping, AAC and AAG were most abundant in pericentric regions of all chromosomes; ACT showed signals on multiple chromosomes at different distances from the centromeres; ATC had its strongest signals in the pericentric region of 4H. A strong signal for AAG in the ONT reads mapped to interstitial regions of 7H in MorexV3 was not observed in the FISH experiments of Cuadrado and Jouve (2007), who worked with cv. Plaisant.

In addition to arrays with short motifs, TRF reported also 26,556 ONT reads with arrays longer than 20 kb and consensus motif lengths above 100 bp. The majority of these (78.6 %) were assigned with high confidence (MAPQ >= 60) to unique positions in the MorexV3 pseudomolecules. The longest array annotated by TRF had 562 copies of a 327 bp motif with high homology to pAS1, spanned 186 kb and mapped 13 Mb away from the distal end of 3HL. We checked whether mapped satellite arrays colocalized with gaps in the MorexV3 pseudomolecules. A scan for tandem repeats in the 1 kb flanking regions of all sequence gaps with TRF found arrays longer than 500 bp in 99 of 147 (65 %) of them. Detected motifs were 2 to 472 bp in size **(Fig. 7)**. Trinucleotides were the most common class; 58 gaps were close to pericentromeric AAG. Apart from trinucleotides, the second most abundant class were 118 bp monomers with high homology to the subtelomeric repeat HvT01. Among four large (> 15 kb) insertions of chloroplast DNA into the nuclear genome, only one coincided with a sequence gap **(Fig. 7)**. These results are consistent with an enrichment of sequence gaps in distal and pericentromeric regions of the pseudomolecules **(Fig. 7, Supplementary Fig. 1-3)**. Taken together, our analyses suggest that the resolution of long low complexity sequences will be the greatest challenge in obtaining T2T assemblies of barley chromosomes.

## Discussion

We have shown that the repeat arrays of telomeres, subtelomeres, centromeres and 5S and 45S rDNA loci are not represented in their entirety in the current barley reference genome sequence assembly (MorexV3). The predominant cause of sequence assembly breakdown is the presence of long, homogeneous tandem repeat arrays that cannot be resolved with reads in the 20 – 100 kb size range and that were also not bridged by contigs of the optical map because DLE-1 label sites were absent from most repeat monomers. By contrast, 45S rDNA was labelled by DLE-1, which enabled us to estimate the abundance of this repeat class in barley. The analysis indicated that regular arrays of the major rDNA loci in chromosomes 5H and 6H spanned over ~21 and ~8 Mb, respectively, and such tandemly organized repetitive sequence cannot be assembled from data obtained by current sequencing and optical mapping technologies whose read lengths exceed 1 Mb only in rare cases. Nevertheless, the presence of marginal parts of the arrays in the assembly, the availability of core rDNA units for each of the major loci and a known quantity of units in each array entertain the possibility of resolving these loci by similar approaches as applied in the T2T assembly of human CHM13 cell line (Nurk et al., 2021).

Our analysis of unassembled repeat sequences cannot explain the discrepancy between assembly size and most flow-cytometric genome size estimates. Even generously rounding up and doubling size estimates for the loci we studied here can account for less than 150 Mb of missing sequence: telomeres 1 Mb; subtelomeres and pericentromeres 60 Mb; ribosomal DNA 40 Mb; and centromeres 40 Mb. However, the difference between the assembly size and the flow-cytometric estimates amounts to 500-800 Mb, i.e. the equivalent of at least an entire barley chromosome. We did not observe in either long- or short-read datasets extended regions with a read depth elevated above the genome-wide average, ruling out the presence of large segmental duplications of low-copy sequence. Doležel et al. (2018) pointed out that their genome size estimates for barley cv. Morex of 4.88 Gbp and 5.04 Gbp, respectively, were obtained assuming a human genome size of 3.257 Gb (GRCh38.p12) and 3.423 Gb (Tiersch et al., 1989), respectively. Recently, Nurk et al. (2021) reported T2T assemblies of all chromosomes in the essentially homozygous human cell line CHM13, amounting to a total assembly size of 3,054,815,472 bp – shorter than either human GSE considered by Doležel et al. (2018). Applying this value yields barley 1C genome size of 4.58 Gb, which is considerably closer to the MorexV3 assembly size and GSEs from long and short reads. The present report focuses on the (as yet incomplete) genome of a single species and cannot provide strong evidence either in favor of or against a revision of flow cytometric size standards. A conclusion in this matter can be drawn only after gapless genome sequences of several plant and animal species have been assembled or if CHM13 could be used as a standard for flow cytometric estimates.

Additional datasets are needed to construct T2T assemblies of barley chromosomes. It may be arguable whether determining copy numbers of all satellite arrays should be prioritized over other research aims addressable by high-throughput sequencing, e.g. expanding the barley pan-genome. However, we are convinced that the complete sequence of a barley centromere would be an important achievement. Near-gapless assembly of the *Arabidopsis thaliana* genome (Naish et al., 2021) enabled epigenomic profiling of centromeres and analysis of transposon insertion patterns. The completion of a centromere of *H. vulgare* would be an important first step towards the comparative sequence and epigenetic analysis of centromere evolution in the genus *Hordeum* and its relation to speciation. Size estimates by us and prior studies (Houben et al., 2007; Presting et al., 1998) indicate that barley centromeres are at least one, possibly three Mb or more in size. Ultra-long ONT reads in the size range of 100 kb – 1 Mb (Prall et al., 2021) might cover a large fraction (5 – 50 %) of a barley centromere. We expect random insertions of *cereba* elements and their subsequent degradation to generate unique patterns in otherwise homogeneous AGAGGG arrays. If reads are long enough to bridge the space between two *cereba* elements, it may be possible to resolve the assembly graphs of barley centromeres into a gapless linear sequence. Another idea is to use epigenetic marks gleaned from long-read sequencing (Gershman et al., 2021) to differentiate between regions identical in DNA sequence.

Knowledge from prior studies using cytological and molecular biology methods has greatly helped in the interpretation of our results. In particular, the near-absence of centromeric sequence might have been hard to ascertain without knowledge of the sequence organization of barley centromeres (Hudakova et al., 2001; Presting et al., 1998). Hi-C contact matrices enabled the precise localization of centromeres, but it would have been difficult to identify *cereba* as a centromere-specific retrotransposon as the majority of *cereba* elements are absent from the sequence assembly. Without FISH mapping of chloroplast probes, we would not have been able to rule out the presence of large chloroplast insertion into the nuclear genomes. This illustrates the importance of complementary methods such as FISH mapping of candidate repeat sequences and ChIP-seq with antibodies for centromeric nucleosomes in assessing assembly completeness.

In this study, we applied an *ad hoc* approach based on BLAST alignment of known motifs to long-reads and *de novo* predictions with TRF to estimate the abundance of tandem repeat arrays. Several methods have been developed to annotate repeats in error-prone long-reads, e.g. the Noise Cancelling Repeat Finder to annotate satellite repeats in noisy long reads (Harris et al., 2019). We deem it a worthwhile subject for future research to develop an analysis toolkit for reference-free repeat prediction and abundance estimation in accurate long-reads similar to the RepeatExplorer2 (Novák et al., 2020) analysis suite for short-read data. Assembly-free repeat analysis from low-coverage (5x) long-read could underpin a more comprehensive assessment of composition and abundance of all classes of tandem repeats, including long satellite arrays that are difficult to study with short reads.

## Methods

### Public datasets used in the study

The MorexV3 assembly is accessible from the European Nucleotide Archive (ENA) under project ID PRJEB40589 and from the Plant Genomics & Phenomics Research Data Repository (PGP, Arend et al. (2016), http://doi.org/10.5447/ipk/2021/3). The ENA accessions for HiFi, ONT and PE450 reads are PRJEB40587, PRJEB40588 and PRJEB31444, respectively. The Bionano optical map is available from PGP (http://doi.org/10.5447/ipk/2021/2). Repeat monomer sequences of HvT01 were downloaded from NCBI (X16095.1:1-118); pAS1 was read from Figure 4 of Brandes et al. (1995). The *cereba* integrase domain was extracted from AY040832.1 based on the sequence shown in Figure 2 of Presting et al. (1998). For 5S rDNA, we used 120 bp (the 5S rRNA gene sequence) from GenBank accession S70723.1.

### Extraction of 5H- and 6H-specific 45S rDNA units

Representative units of 5H and 6H rDNA loci were extracted from unassigned contigs of MorexV3 assembly using a two-step procedure. First, 86 and 48 complete but heterogenous rDNA units found in marginal parts of 45S rDNA arrays present in 5H and 6H pseudomolecule, respectively, were used to construct consensual 5H and 6H units. The chromosome-specific consensuses were then applied for BLAST search in unassigned contigs (chrUn) of the MorexV3 presumed to harbor collapsed homogenous units forming cores of the major rDNA arrays. We identified 264 and 25 identical rDNA monomers for the 5H and 6H variants, respectively, which we used as representative units for the 5H and 6H loci.

### Read mapping to MorexV3 and genome size estimation with k-mers

HiFi, ONT and PE450 were aligned to the MorexV3 pseudomolecules with Minimap2 version 2.17 (Li, 2018) using the presets map-pb, map-ont and sr, respectively. PE450 were processed with cutadapt (Martin, 2011) prior to alignment. Alignment records were converted to Binary Alignment Map (BAM) format using SAMtools (Li et al., 2009) and sorted with Novosort (http://www.novocraft.com/products/novosort/). Read depth was calculated with SAMtools and aggregated in 1 kb windows with BEDtools (Quinlan and Hall, 2010). Summary statistics and GSEs were calculated in R (R Core Team, 2017). Genome size estimation based on *k*-mer spectra was done for HiFi, ONT and PE40 reads with findGSE (Sun et al., 2018) using Jellyfish (Marçais and Kingsford, 2011) for k-mer counting.

### Tandem repeat annotation and quantification in long-reads

Tandem repeats were identified with Tandem Repeat Finder (TRF, Benson (1999)) using the parameter setting “2 5 7 80 10 50 2000 -l 1 -h”. Read files in FASTQ format were converted to FASTA format with seqtk (https://github.com/lh3/seqtk). GNU Parallel (Tange, 2018) was used to process reads in parallel. TRF was run on HiFi reads, ONT reads, the MorexV3 pseudomolecules, and 1 kb flanking regions of MorexV3 gaps. Prior to further analysis, detected motifs were converted to a canonical form, namely the lexically minimal sequence among all cyclic shifts of the motif and its reverse complement. For example, GAG is synonymous with CTC, CCT, TCC, GGA and AGG; AGG is the canonical motif. Summary statistics were calculated and plots were generated using functions of the R statistical environment (R Core Team, 2017).

### Quantification of repeats in long-reads by sequence alignment

Monomer sequences of repeat arrays were aligned to different references using BLASTN (Altschul et al., 1990) with default parameters (BLAST+, version 2.2.30). The references were: the MorexV3 pseudomolecules, HiFi reads, ONT reads and 1 kb flanking regions of gaps in MorexV3. Overlapping alignments were merged with BEDTools (Quinlan and Hall, 2010). Statistical analysis was done in R (R Core Team, 2017).

### Quantification of 45S ribosomal DNA with Bionano map data

Regular arrays of 45S rDNA units can be recognized and quantified in optical maps generated on the Saphyr platform (Bionano Genomics, San Diego, USA) thanks to the presence of a DLE-1 labelled site in the 26S rRNA gene, which generates a regular pattern with ~9 - 10 kb label spacing (**Figure 6a**). The units were quantified from size-filtered (>150 kb) raw (single-molecule) data of DLE-1 optical map of barley cv. Morex (Mascher et al. 2021), applying a RefAligner (Bionano Genomics) function simpleRepeatStandalone and repeat stretch tolerance of 0.1. Arrays of six and more repeat units were considered. Unit size estimates obtained from the optical map data were corrected using the coefficient of 0.952 to eliminate error due to 4.8 % expansion of the optical map compared to the sequence, calculated from a sequence-to-map alignment. The resulting rmap file was analyzed in Microsoft Excel and unit size and number of units were plotted in a histogram for visual analysis.

### ChIP-seq

Nuclei were isolated as described previously (Neumann et al., 2012) from *Hordeum vulgare* cv. Morex 4 days-germinated embryos. ChIP-seq protocol from the same publication was followed with minor modifications using the anti-barley α-CENH3 antibody (Sanei et al., 2011). Briefly, nuclei isolated from 4 g tissue were centrifuged at 600 g for 15 min at 4°C and resuspended in 1 ml micrococcal nuclease (MNase) buffer (10% sucrose, 50 mM Tris-HCl pH 7.5, 4 mM MgCl_2_, 1 mM CaCl2, 1× protease inhibitor cocktail (cOmplete™ Roche)), divided into 10 aliquots and digested with the range of MNase amounts (NEB M0247S, 500-2000 GU of the enzyme per aliquot) for 10 min at 37°C, yielding fragments within the range between mono- and tetra-nucleosomal size. The reactions were stopped by adding 0.5 M EDTA to a final concentration of 20 mM and samples were pooled and centrifuged at 13,000 g for 5 min at 4°C. The supernatant containing well-digested chromatin was saved while the pellet containing poorly digested chromatin was redigested with 500 units of MNase for 5 min at 37°C in 200 μl MNase buffer. The reaction was stopped with EDTA and centrifuged as described above. The chromatin fractions were pooled, resulting in >75% consisting of mononucleosomes, and diluted with the same volume of ChIP incubation buffer (20 mM Tris pH 7.5, 140 mM NaCl, 1 mM EDTA pH8, 1% Triton X-100, 0.1% sodium deoxycholate, 0.1% sodium dodecyl sulfate (SDS), 1× protease inhibitor). A 50 μl aliquot was taken for DNA isolation to serve as an input control sample. Antibody binding was done for 2 h at 4°C in 200 μl of PBS buffer with 0.01% Tween-20 containing 30 μl magnetic beads and 9 μg of the antibody. The beads with bound antibody were mixed with the chromatin and the mixture was incubated with rotation overnight at 4°C. Immunoprecipitated complexes were washed 3×5 min using 800 μl of the ChIP incubation buffer, followed by two washes with the ChIP incubation buffer containing 300mM NaCl and two washes with TE buffer. Elution of the chromatin was done using 2×100 μl of elution buffer (1 % SDS in TE with proteinase K) for 15 min at 55°C. DNA from the ChIP and input control samples was isolated using ChIP DNA Clean and Concentrator Kit (Zymo Research, Irvine, CA) to prepare sequencing libraries using NEBnext Ultra II kit (NEB). Paired-end sequencing was done on S1 flow-cell using NovaSeq.

### ChIP-seq data analysis

Raw ChIP-seq reads were trimmed to 120 bp, and adapters and low-quality reads were removed by TrimGalore (https://github.com/FelixKrueger/TrimGalore). Trimmed reads were mapped to the MorexV3 reference with Minimap2 (Li, 2018). Alignment records were converted to BAM format with SAMtools (Li et al., 2009) and sorted and deduplicated with Novosort (http://www.novocraft.com/products/novosort/). The counts of uniquely mapped, non-duplicated reads (samtools view -q 20 -F 3332) were aggregated in 1 kb windows for visualization. The enrichment of repetitive sequences in the ChIP-seq data was evaluated using RepeatExplorer2 followed by ChIP-seq Mapper, both integrated at the Galaxy server (https://repeatexplorer-elixir.cerit-sc.cz/galaxy/). First, the repetitive DNA cluster database was generated by RepeatExplorer2 using similarity-based clustering of 2×100 bp WGS Illumina reads (SRA accession ERR125905). ChIP and Input reads were then mapped to contigs resulting from the clustering, and the CENH3-enriched clusters were determined based on ChIP/Input reads elevated ratio.

### Fluorescence in situ hybridization (FISH)

Chromosome spreads of barley cv. Morex was prepared as described by Aliyeva-Schnorr et al. (2015). The barley cpDNA-bearing BAC clone ChHB040G01 (Martis et al., 2012), subtelomeric satellite repeat HvT01 (Belostotsky and Ananiev, 1990b) and 45S rDNA containing clone pTa71 (Gerlach and Bedbrook, 1979) were labelled with dUTP-ATTO550, dUTP-ATTO488 and dUTP-ATTO647, respectively, as FISH probes using nick translation labelling kits (Jena Bioscience). Before hybridization, slides were treated with 45% acetic acid at RT for 10 min, followed by 0.1% pepsin/ 0.01 N HCl at 37°C for 10 min and post-fixed in 4% paraformaldehyde nat RT for 10 min. The hybridization mixture contained 50% (v/v) formamide, 10% (w/v) dextran sulfate, 2× SSC and 5 ng/μl of each probe. Chromosomal DNAs and probes were denatured at 75°C for 2 min, followed by hybridization at 37°C for 20-24 hr. The final stringent wash was performed in 2× SSC at 57°C for 20 min and slides were dehydrated in 70-90-100% ethanol series for 3 min each. Chromosomes were counterstained by 10 μg/ml 4Ͱ,6-diamidino-2-phenylindole (DAPI) in Vectashield antifade mounting medium (Vector Laboratories). Images were captured using an epifluorescence microscope BX61 (Olympus) equipped with a cooled CCD camera (Orca ER, Hamamatsu) and pseudocolored using Adobe Photoshop CS6.

## Accession codes

CENH3 ChIP-seq data are accessible from the European Nucleotide Archive (ENA, https://www.ebi.ac.uk/ena) under project ID PRJEBXXX.

## Author contributions

MM, HŠ, NS, AH, JD conceived the study. HT, ZT and HŠ collected and analyzed optical mapping data. PN performed ChIP-seq experiments. PN and MM analyzed sequence data. AH contributed CENH3 antibodies. YTK performed FISH. MM, PN and HŠ wrote the paper with input from all co-authors.

## Acknowledgments

This research was supported by the grant MITOCHROM jointly funded by the German Research Foundation (DFG, grant MA 6611/4-1 to M.M.) and the Czech Science Foundation (GAČR, grant 18-14450J to J.D.). Further support was provided by the German Ministry of Education and Research (BMBF) in frame of the grants SHAPE II (FKZ 031B0884 to N.S. and M.M.) and by GAČR (grant 17-17564S to H.Š.) We sincerely thank Petr Novák from The Institute of Plant Molecular Biology, Biology Centre CAS for valuable discussions on RepeatExplorer data analysis. Computational resources were provided by the project “e-Infrastruktura CZ” (e-INFRA LM2018140) provided within the program Projects of Large Research, Development and Innovations Infrastructures and ELIXIR-CZ project (LM2018131), part of the international ELIXIR infrastructure.

## Supplementary Items

**Supplementary Figure 1:** Read depth in HiFi reads.

**Supplementary Figure 2:** Read depth in ONT reads.

**Supplementary Figure 3:** Read depth in PE450 reads.

**Supplementary Figure 4:** Sequence organization of the pericentromeric region of chromosome 2H.

**Supplementary Figure 5:** Sequence organization of the pericentromeric region of chromosome 3H.

**Supplementary Figure 6:** Sequence organization of the pericentromeric region of chromosome 4H.

**Supplementary Figure 7:** Sequence organization of the pericentromeric region of chromosome 5H.

**Supplementary Figure 8:** Sequence organization of the pericentromeric region of chromosome 6H.

**Supplementary Figure 9:** Sequence organization of the pericentromeric region of chromosome 7H.

**Supplementary Figure 10:** (Peri)centromeric region of chromosome 2H in the optical map.

**Supplementary Figure 11:** α-CENH3-ChIP-Seq Mapper analysis.

**Supplementary Figure 12**. 45S ribosomal DNA in Morex V3 assembly.

