## Supplementary Figures for "Prospects of telomere-to-telomere assembly in barley: analysis of sequence gaps in the MorexV3 reference genome"

### Slide 1
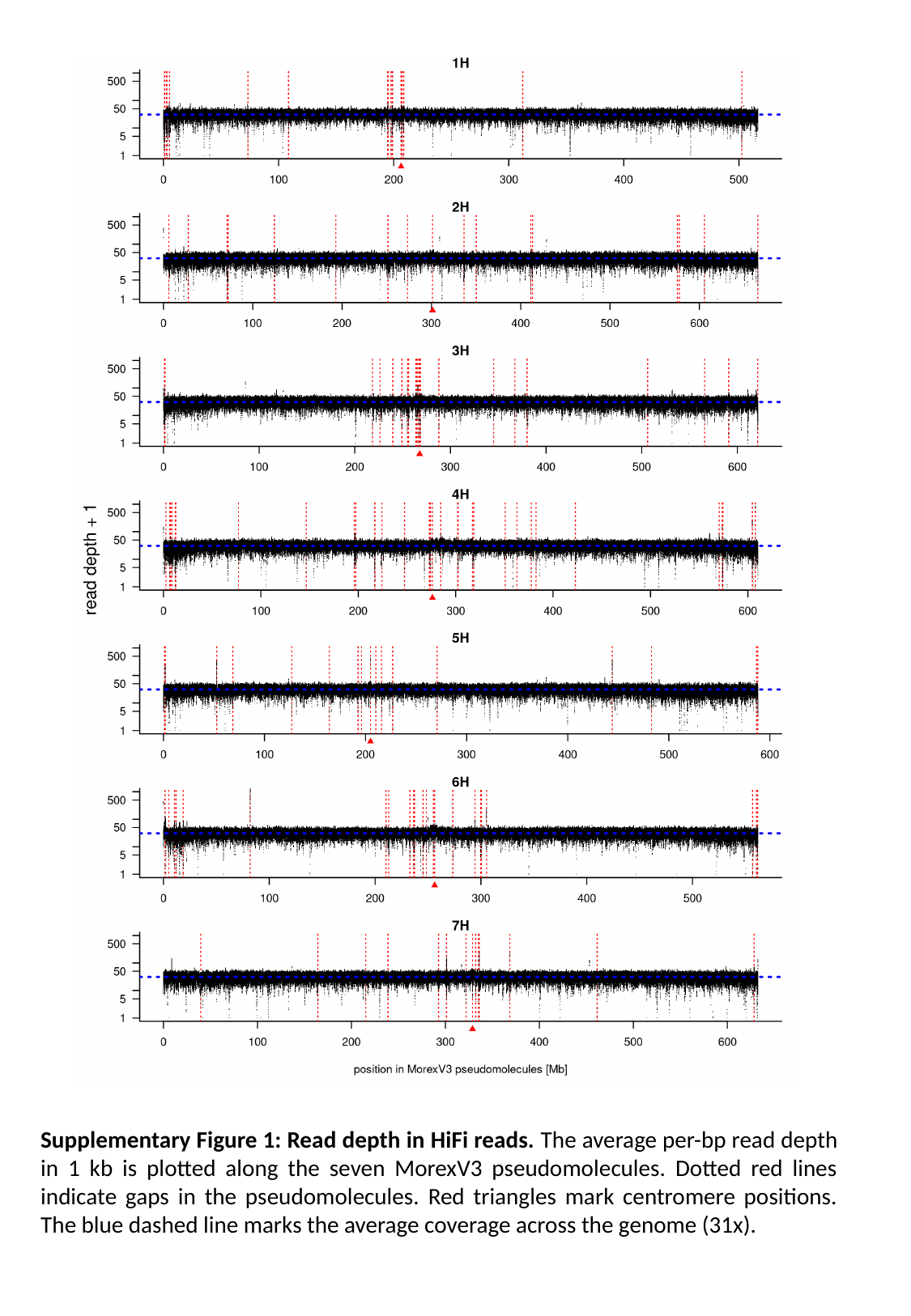

Supplementary Figure 1: Read depth in HiFi reads. The average per-bp read depth in 1 kb is plotted along the seven MorexV3 pseudomolecules. Dotted red lines indicate gaps in the pseudomolecules. Red triangles mark centromere positions. The blue dashed line marks the average coverage across the genome (31x).

### Slide 2
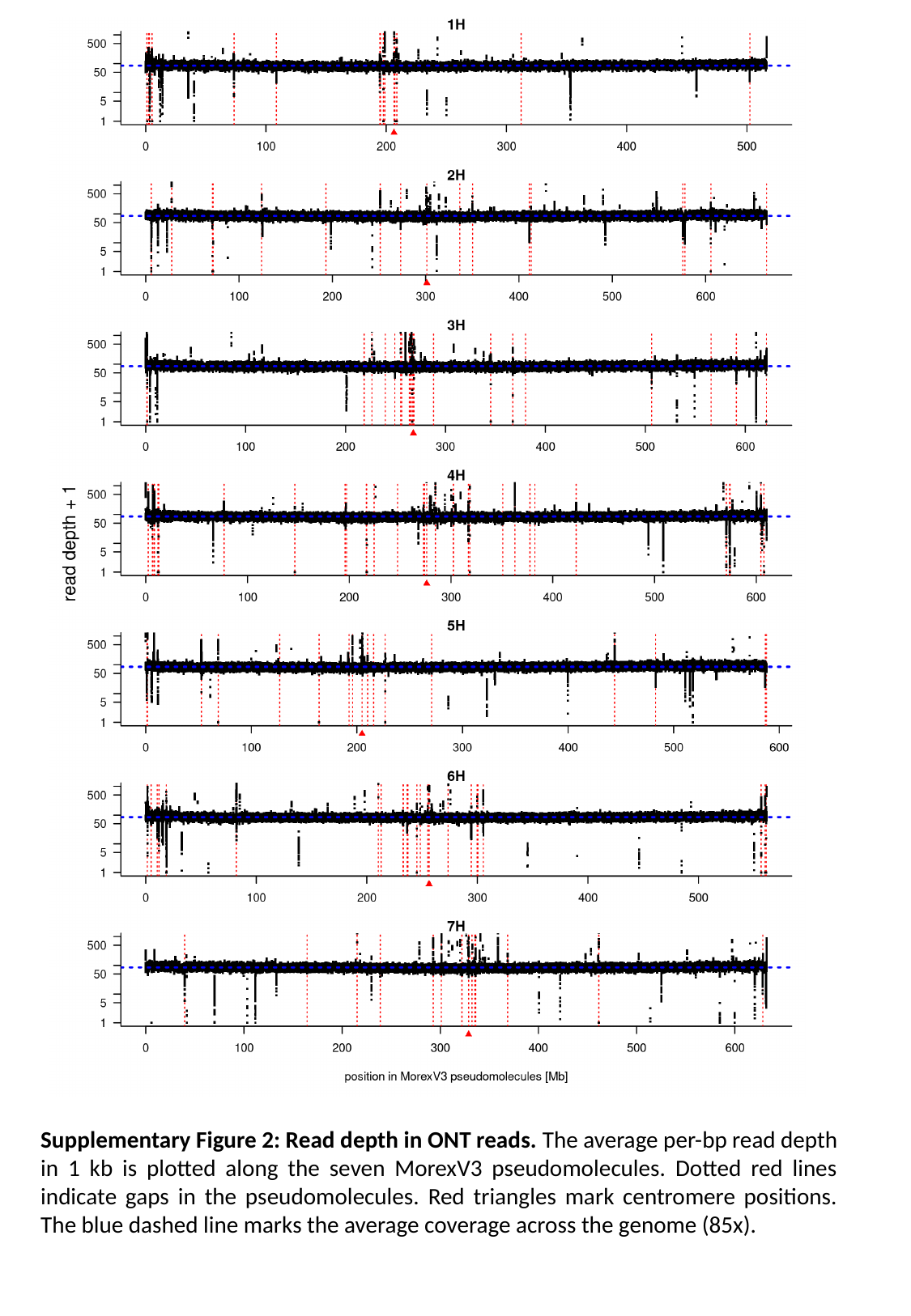

Supplementary Figure 2: Read depth in ONT reads. The average per-bp read depth in 1 kb is plotted along the seven MorexV3 pseudomolecules. Dotted red lines indicate gaps in the pseudomolecules. Red triangles mark centromere positions. The blue dashed line marks the average coverage across the genome (85x).

### Slide 3
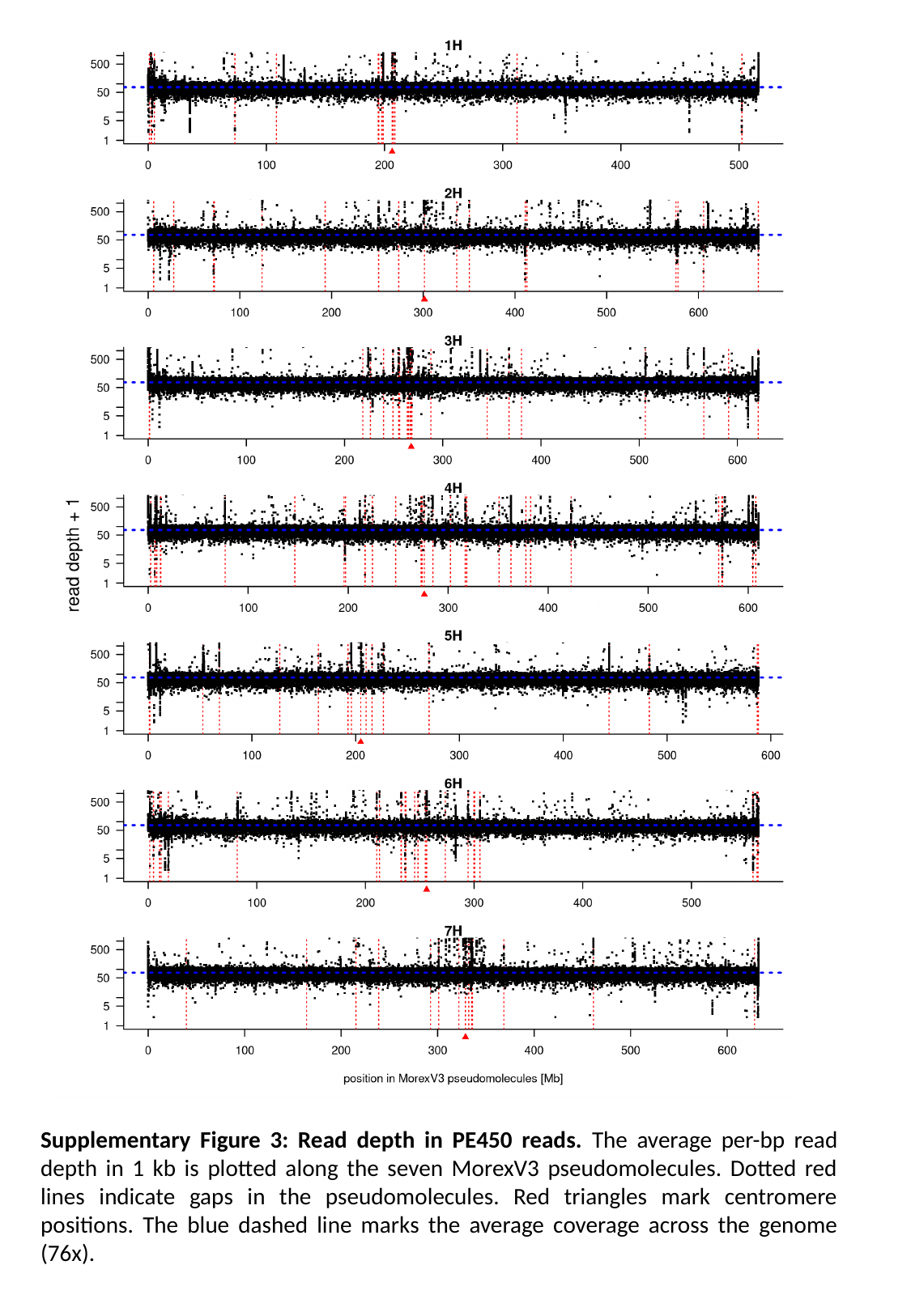

Supplementary Figure 3: Read depth in PE450 reads. The average per-bp read depth in 1 kb is plotted along the seven MorexV3 pseudomolecules. Dotted red lines indicate gaps in the pseudomolecules. Red triangles mark centromere positions. The blue dashed line marks the average coverage across the genome (76x).

### Slide 4
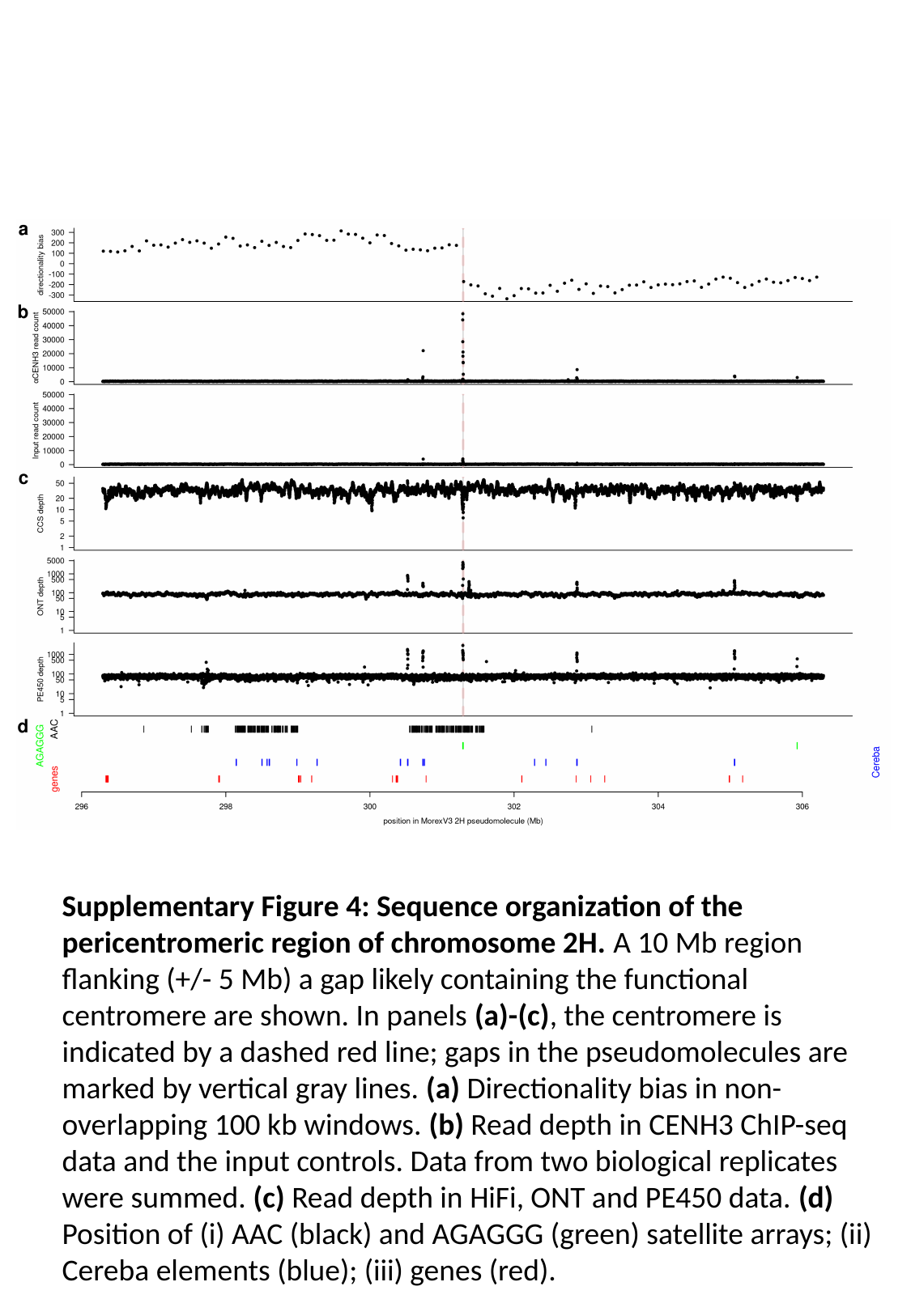

Supplementary Figure 4: Sequence organization of the pericentromeric region of chromosome 2H. A 10 Mb region flanking (+/- 5 Mb) a gap likely containing the functional centromere are shown. In panels (a)-(c), the centromere is indicated by a dashed red line; gaps in the pseudomolecules are marked by vertical gray lines. (a) Directionality bias in non-overlapping 100 kb windows. (b) Read depth in CENH3 ChIP-seq data and the input controls. Data from two biological replicates were summed. (c) Read depth in HiFi, ONT and PE450 data. (d) Position of (i) AAC (black) and AGAGGG (green) satellite arrays; (ii) Cereba elements (blue); (iii) genes (red).

### Slide 5
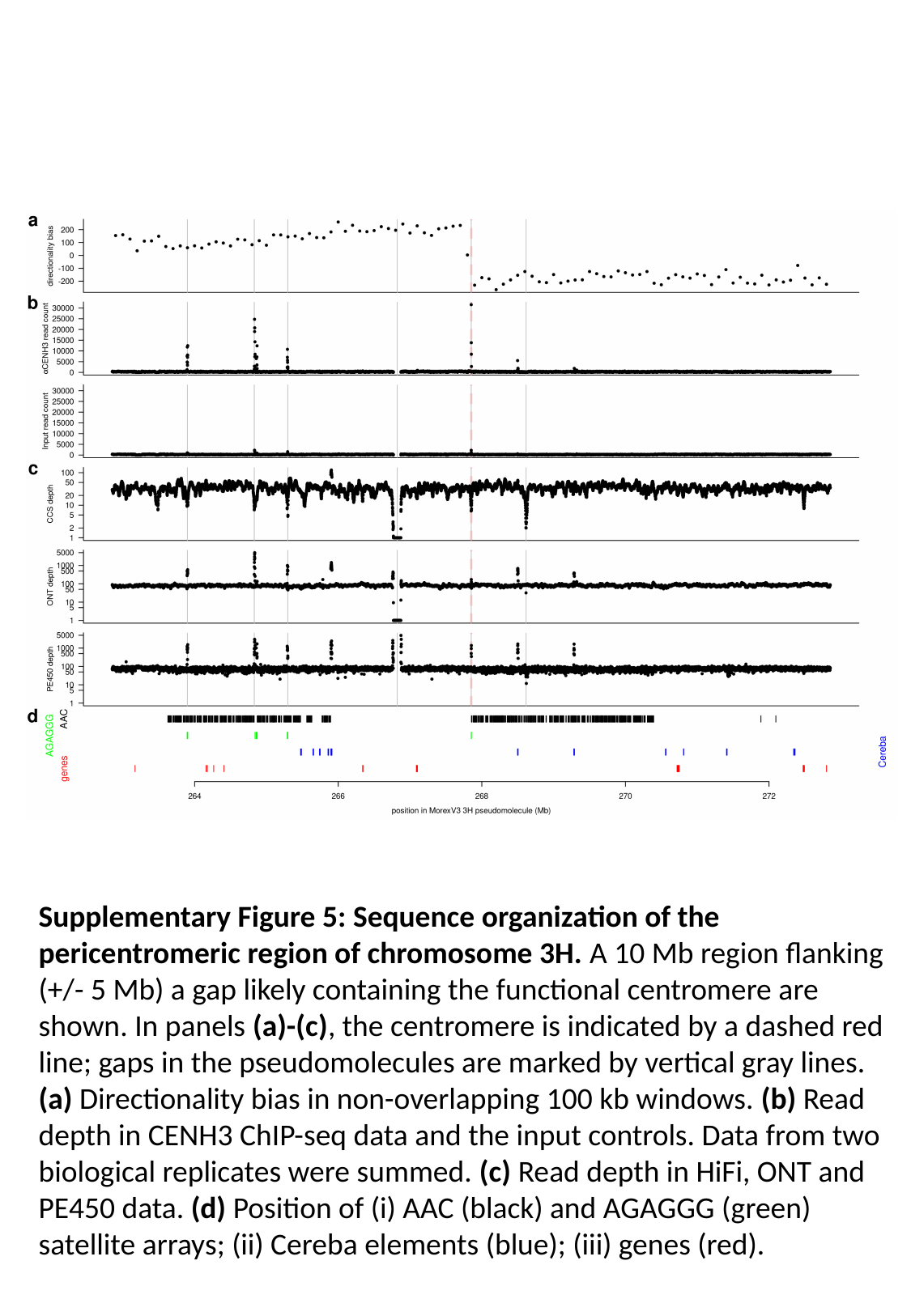

Supplementary Figure 5: Sequence organization of the pericentromeric region of chromosome 3H. A 10 Mb region flanking (+/- 5 Mb) a gap likely containing the functional centromere are shown. In panels (a)-(c), the centromere is indicated by a dashed red line; gaps in the pseudomolecules are marked by vertical gray lines. (a) Directionality bias in non-overlapping 100 kb windows. (b) Read depth in CENH3 ChIP-seq data and the input controls. Data from two biological replicates were summed. (c) Read depth in HiFi, ONT and PE450 data. (d) Position of (i) AAC (black) and AGAGGG (green) satellite arrays; (ii) Cereba elements (blue); (iii) genes (red).

### Slide 6
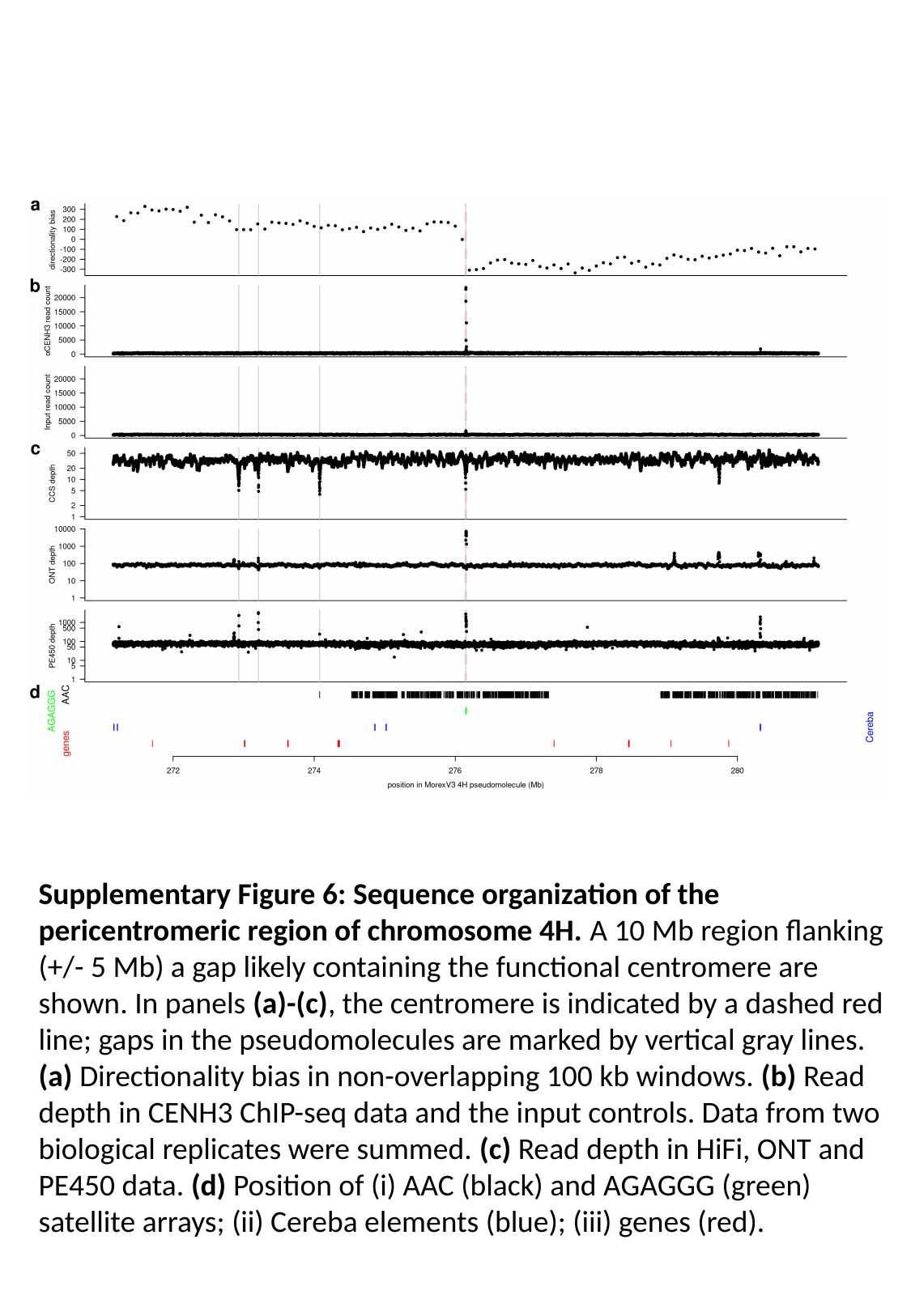

Supplementary Figure 6: Sequence organization of the pericentromeric region of chromosome 4H. A 10 Mb region flanking (+/- 5 Mb) a gap likely containing the functional centromere are shown. In panels (a)-(c), the centromere is indicated by a dashed red line; gaps in the pseudomolecules are marked by vertical gray lines. (a) Directionality bias in non-overlapping 100 kb windows. (b) Read depth in CENH3 ChIP-seq data and the input controls. Data from two biological replicates were summed. (c) Read depth in HiFi, ONT and PE450 data. (d) Position of (i) AAC (black) and AGAGGG (green) satellite arrays; (ii) Cereba elements (blue); (iii) genes (red).

### Slide 7
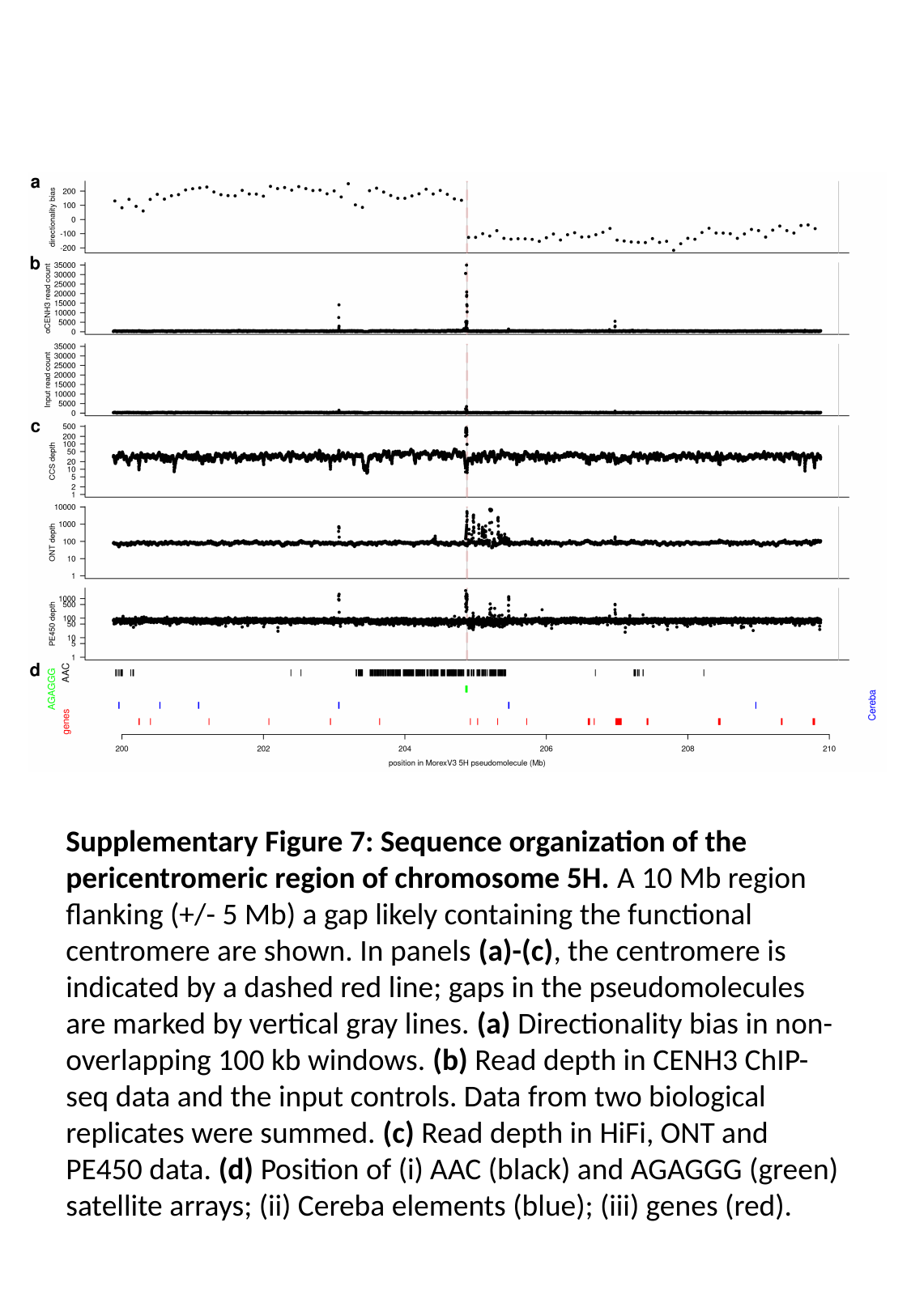

Supplementary Figure 7: Sequence organization of the pericentromeric region of chromosome 5H. A 10 Mb region flanking (+/- 5 Mb) a gap likely containing the functional centromere are shown. In panels (a)-(c), the centromere is indicated by a dashed red line; gaps in the pseudomolecules are marked by vertical gray lines. (a) Directionality bias in non-overlapping 100 kb windows. (b) Read depth in CENH3 ChIP-seq data and the input controls. Data from two biological replicates were summed. (c) Read depth in HiFi, ONT and PE450 data. (d) Position of (i) AAC (black) and AGAGGG (green) satellite arrays; (ii) Cereba elements (blue); (iii) genes (red).

### Slide 8
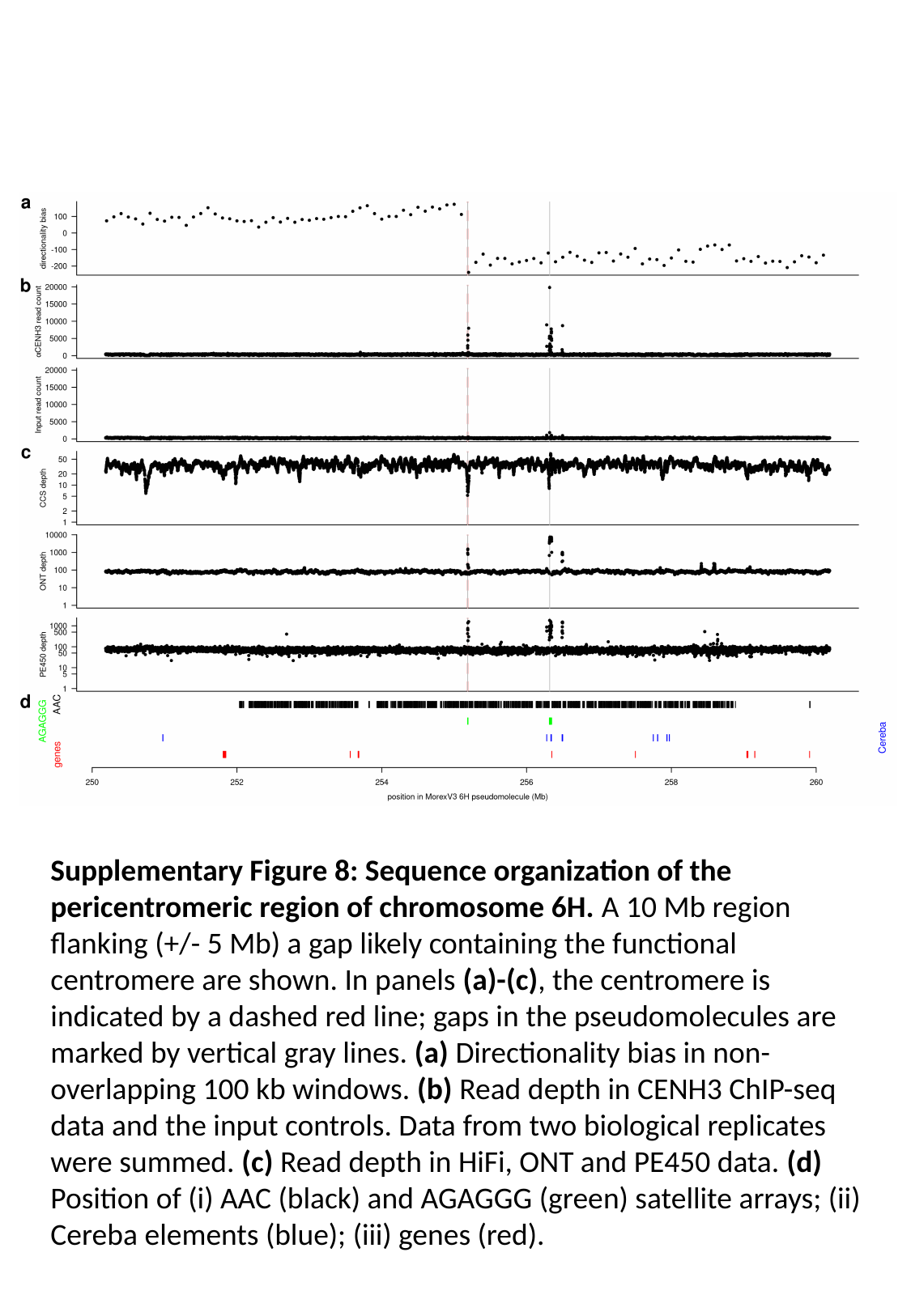

Supplementary Figure 8: Sequence organization of the pericentromeric region of chromosome 6H. A 10 Mb region flanking (+/- 5 Mb) a gap likely containing the functional centromere are shown. In panels (a)-(c), the centromere is indicated by a dashed red line; gaps in the pseudomolecules are marked by vertical gray lines. (a) Directionality bias in non-overlapping 100 kb windows. (b) Read depth in CENH3 ChIP-seq data and the input controls. Data from two biological replicates were summed. (c) Read depth in HiFi, ONT and PE450 data. (d) Position of (i) AAC (black) and AGAGGG (green) satellite arrays; (ii) Cereba elements (blue); (iii) genes (red).

### Slide 9
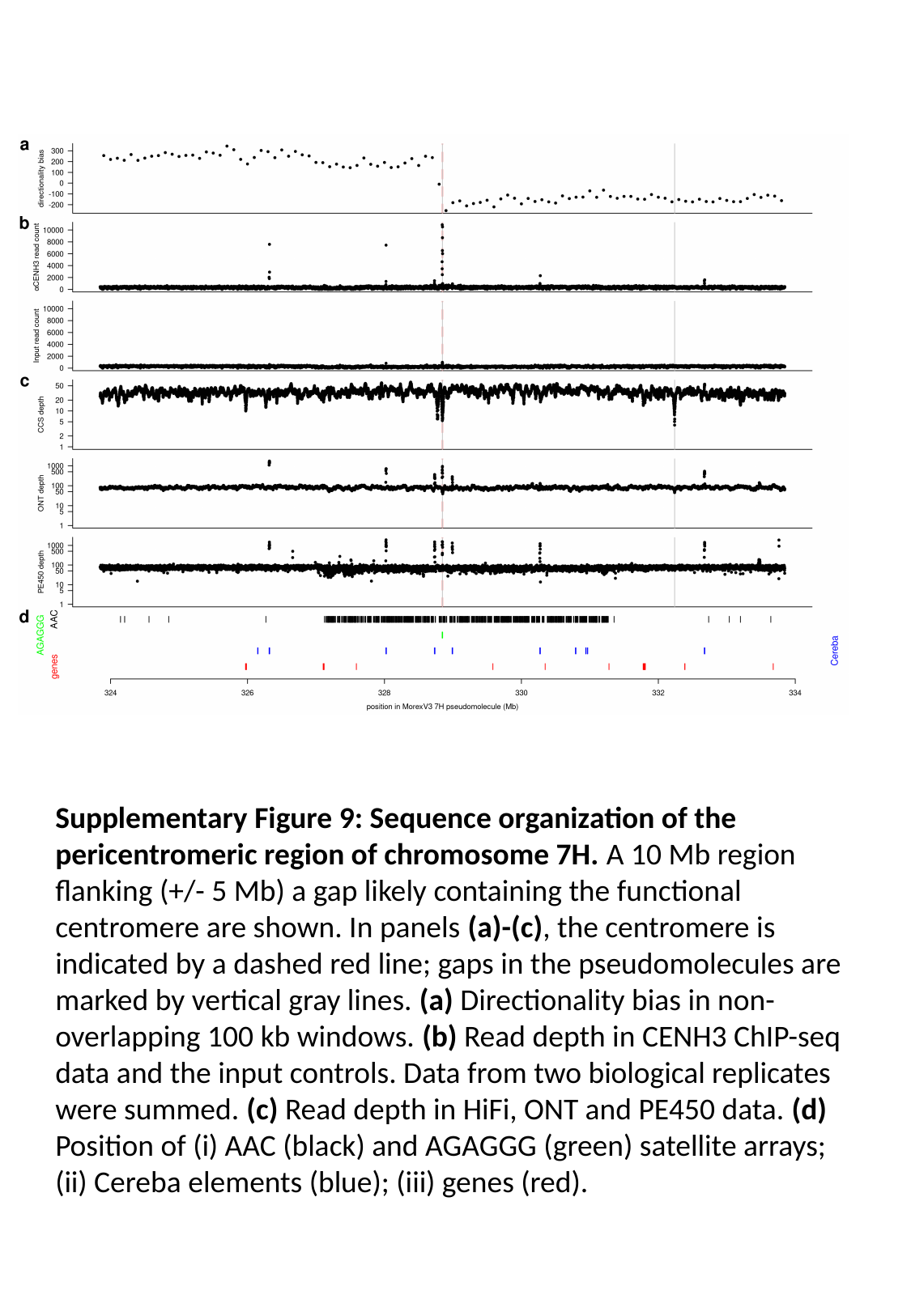

Supplementary Figure 9: Sequence organization of the pericentromeric region of chromosome 7H. A 10 Mb region flanking (+/- 5 Mb) a gap likely containing the functional centromere are shown. In panels (a)-(c), the centromere is indicated by a dashed red line; gaps in the pseudomolecules are marked by vertical gray lines. (a) Directionality bias in non-overlapping 100 kb windows. (b) Read depth in CENH3 ChIP-seq data and the input controls. Data from two biological replicates were summed. (c) Read depth in HiFi, ONT and PE450 data. (d) Position of (i) AAC (black) and AGAGGG (green) satellite arrays; (ii) Cereba elements (blue); (iii) genes (red).

### Slide 10
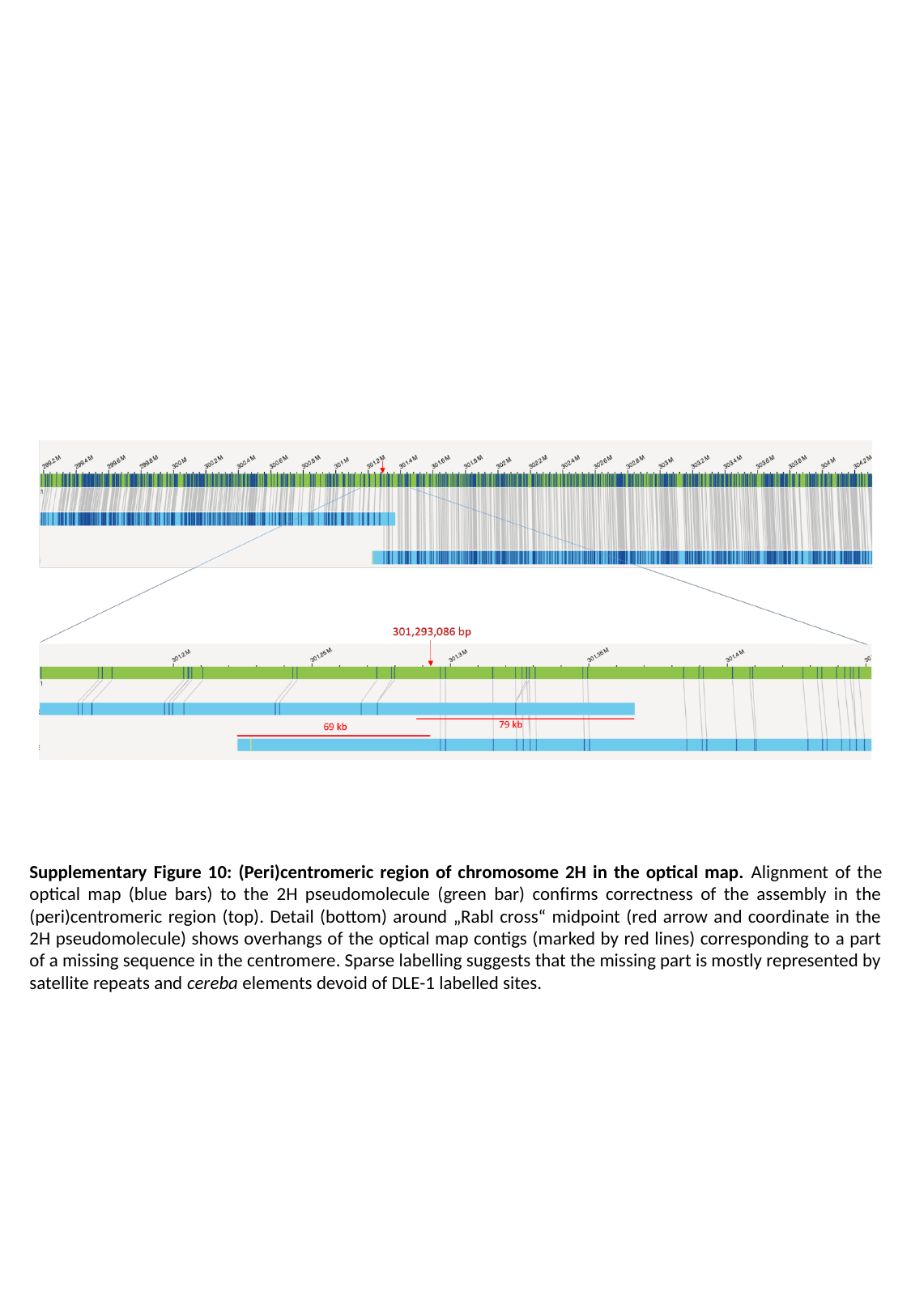

Supplementary Figure 10: (Peri)centromeric region of chromosome 2H in the optical map. Alignment of the optical map (blue bars) to the 2H pseudomolecule (green bar) confirms correctness of the assembly in the (peri)centromeric region (top). Detail (bottom) around „Rabl cross“ midpoint (red arrow and coordinate in the 2H pseudomolecule) shows overhangs of the optical map contigs (marked by red lines) corresponding to a part of a missing sequence in the centromere. Sparse labelling suggests that the missing part is mostly represented by satellite repeats and cereba elements devoid of DLE-1 labelled sites.

### Slide 11
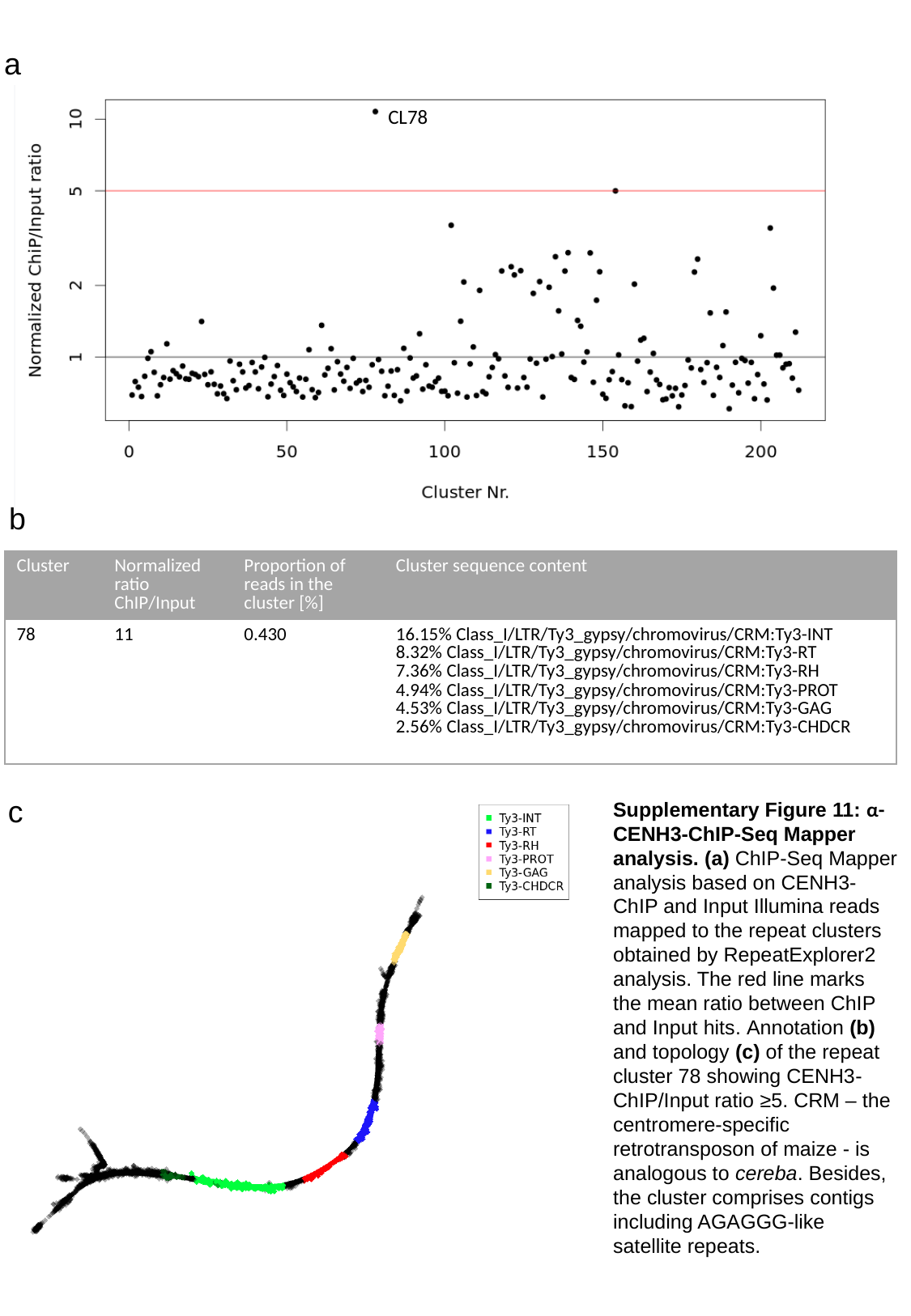

a
CL78
b
| Cluster | Normalized ratio ChIP/Input | Proportion of reads in the cluster [%] | Cluster sequence content |
| --- | --- | --- | --- |
| 78 | 11 | 0.430 | 16.15% Class\_I/LTR/Ty3\_gypsy/chromovirus/CRM:Ty3-INT 8.32% Class\_I/LTR/Ty3\_gypsy/chromovirus/CRM:Ty3-RT 7.36% Class\_I/LTR/Ty3\_gypsy/chromovirus/CRM:Ty3-RH 4.94% Class\_I/LTR/Ty3\_gypsy/chromovirus/CRM:Ty3-PROT 4.53% Class\_I/LTR/Ty3\_gypsy/chromovirus/CRM:Ty3-GAG 2.56% Class\_I/LTR/Ty3\_gypsy/chromovirus/CRM:Ty3-CHDCR |
c
Supplementary Figure 11: α-CENH3-ChIP-Seq Mapper analysis. (a) ChIP-Seq Mapper analysis based on CENH3-ChIP and Input Illumina reads mapped to the repeat clusters obtained by RepeatExplorer2 analysis. The red line marks the mean ratio between ChIP and Input hits. Annotation (b) and topology (c) of the repeat cluster 78 showing CENH3-ChIP/Input ratio ≥5. CRM – the centromere-specific retrotransposon of maize - is analogous to cereba. Besides, the cluster comprises contigs including AGAGGG-like satellite repeats.

### Slide 12
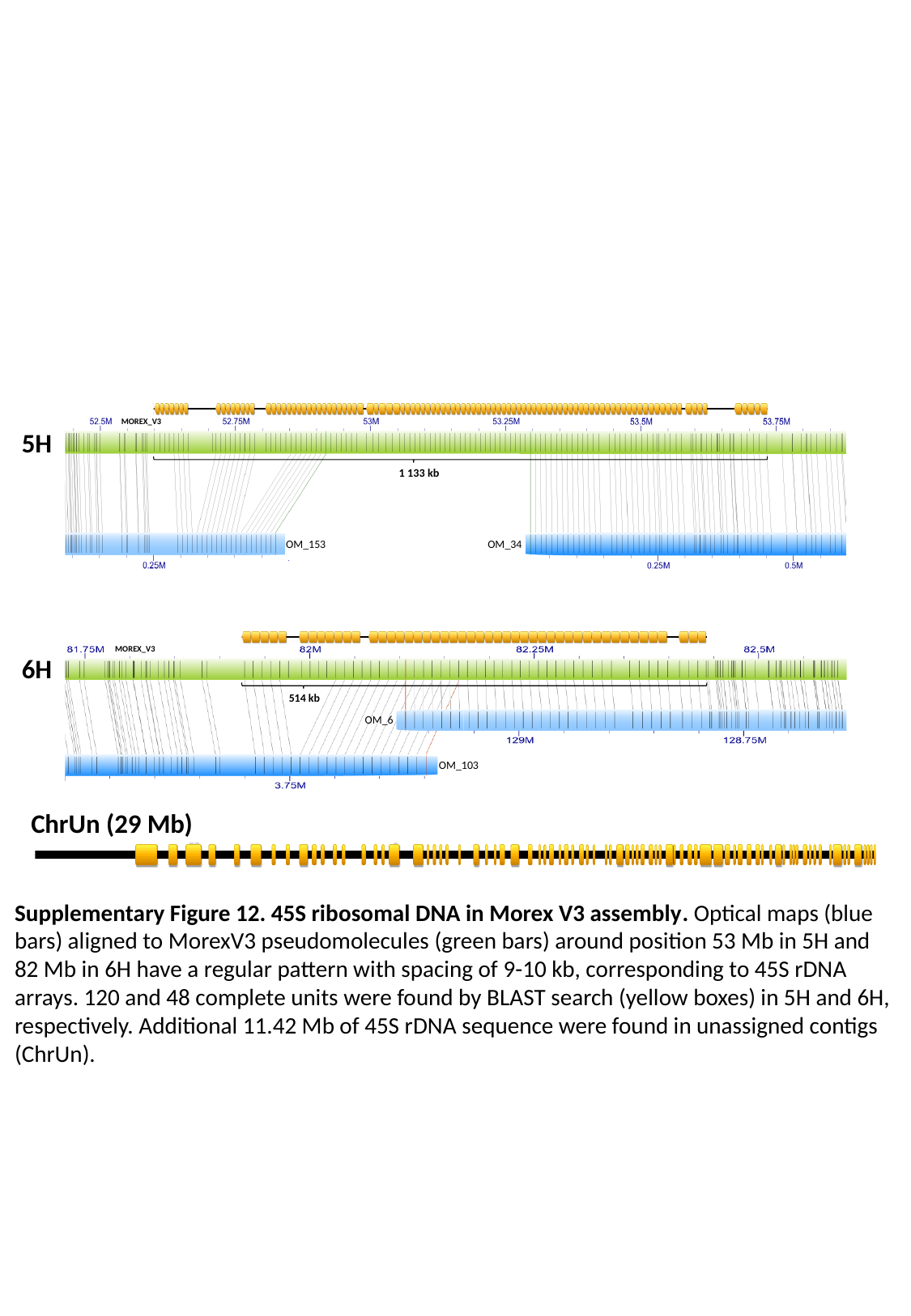

MOREX_V3
5H
1 133 kb
OM_153
OM_34
MOREX_V3
6H
514 kb
OM_6
OM_103
ChrUn (29 Mb)
Supplementary Figure 12. 45S ribosomal DNA in Morex V3 assembly. Optical maps (blue bars) aligned to MorexV3 pseudomolecules (green bars) around position 53 Mb in 5H and 82 Mb in 6H have a regular pattern with spacing of 9-10 kb, corresponding to 45S rDNA arrays. 120 and 48 complete units were found by BLAST search (yellow boxes) in 5H and 6H, respectively. Additional 11.42 Mb of 45S rDNA sequence were found in unassigned contigs (ChrUn).
Bionano compute assembly
